# Suppression of epileptic seizures by transcranial activation of K^+^-selective channelrhodopsin

**DOI:** 10.1101/2024.01.03.573747

**Authors:** Xiaodong Duan, Chong Zhang, Yujie Wu, Jun Ju, Zhe Xu, Xuanyi Li, Yao Liu, Schugofa Ohdah, Oana M. Constantin, Zhonghua Lu, Cheng Wang, Xiaojing Chen, Christine E. Gee, Georg Nagel, Sheng-Tao Hou, Shiqiang Gao, Kun Song

## Abstract

Optogenetics is a valuable tool for studying the mechanisms of neurological diseases and is now being developed for therapeutic applications. In rodents and macaques, improved channelrhodopsins have been applied to achieve transcranial optogenetic stimulation. While transcranial photoexcitation of neurons has been achieved, noninvasive optogenetic inhibition for treating hyperexcitability-induced neurological disorders has remained elusive. There is a critical need for effective inhibitory optogenetic tools that are highly light-sensitive and capable of suppressing neuronal activity in deep brain tissue. In this study, we developed a highly sensitive K^+^-conductive channelrhodopsin (hsKCR) by molecular engineering of the recently discovered *Hyphochytrium catenoides* kalium (potassium) channelrhodopsin 1. Transcranial activation of hsKCR significantly prolongs the time to the first seizure, increases survival, and decreases seizure activity in several mouse epileptic models. Our approach for transcranial optogenetic inhibition of neural hyperactivity may be adapted for cell type-specific neuromodulation in both basic and preclinical settings.

## Introduction

Many neuromodulatory strategies using optogenetics have been developed and applied to treat neurological diseases in translational settings^1,2^. For example, promising results have emerged from clinical trials using channelrhodopsins (ChRs) to treat retinitis pigmentosa-related blindness^3^. Optogenetics also enables targeted neural stimulation to treat brain disorders in animal models, such as epilepsy, addiction, and Parkinson’s disease^4-6^. However, a significant drawback of applying optogenetics to the brain is the requirement for surgery and invasive implantation of hardware for delivering light into deep neural tissues. This carries the risk of permanent brain injury and infection^7^. Recent advancements in deep transcranial optogenetics have been made to avoid these risks. Tools like ChRmine and SOUL allow transcranial optogenetic excitation of neural activity up to depths of 5-7 mm in animal behavioral paradigms^8,9^. While transcranial neural excitation has been achieved, optical inhibition has proven challenging due to a lack of ultra-sensitive inhibitory optogenetic tools suitable for deep brain silencing^10^. Light-driven proton pumps (e.g., archaerhodopsin: eArch3.0), chloride pumps (e.g., halorhodopsin: eNpHR3.0/Jaws) and light-gated anion-conducting channelrhodopsins (ACRs) are established tools for inhibiting neuronal activity^11-14^. Nevertheless, prolonged illumination of these tools can disrupt intracellular pH or change the reversal potential of GABAA receptors, which may result in neural activation rather than inhibition^15,16^. Thus, better inhibitory optogenetic tools should be developed.

Given the essential role of K^+^ conductance in the termination of action potentials, the activation of K^+^ channels holds promise as a therapeutic approach for diseases characterized by neuronal hyperexcitability, for example epilepsy, stroke and Alzheimer’s disease^17-20^. Considerable interest and efforts have therefore been devoted to the development of engineered light-gated K^+^-selective channels for optical inhibition. Previous studies have explored various strategies, such as the fusion of the miniature viral potassium channel Kcv with a blue light-sensitive LOV2 module to create BLINKs^21^, and the use of a two-component optical silencer system (PAC-K) consisting of photoactivated adenylyl cyclase (PAC) and the cyclic nucleotide-gated potassium channel SthK^22^. Although synthetic light-gated potassium channels can efficiently inhibit neural activity, these approaches have certain limitations that hinder their widespread application *in vivo*. For instance, they typically respond to a relatively narrow range of blue light, exhibit slow channel kinetics, have large gene sizes, show irreversible activation, and display poor expression in mammalian cells^23,24^. These drawbacks restrict their utility in living animals.

Here, we utilized the recently discovered K^+^-selective channelrhodopsins (KCRs) as a starting point and engineered a K^+^-selective ChR with significantly enhanced photocurrents and light sensitivity^25^. The new KCR variant was named hsKCR for <u>h</u>ighly <u>s</u>ensitive <u>k</u>alium (K^+^) <u>c</u>hannel<u>r</u>hodopsin. Since its activation is ultra-sensitive to light, it enables noninvasive optogenetic neural silencing via illuminating through the intact skull. Using a well-established drug-induced acute seizure model, we tested its inhibitory effects *in vivo* and observed that green-light stimulation of hsKCR suppressed kainate (KA)-induced epileptic discharges and significantly reduced behavioral signs of epileptic seizures. To further assess the therapeutic potential of hsKCR, we developed a bilateral transcranial optogenetic (BTO) strategy based on hsKCR and evaluated its seizure suppression capabilities in different seizure models. Our results demonstrated that the hsKCR-based BTO approach effectively inhibited pilocarpine-induced neural activity, controlled KA-induced seizures, and significantly decreased seizures and improved survival of pentylenetetrazol (PTZ)-induced epileptic mice.

## Results

### Engineering and characterization of a highly sensitive light-gated K^+^ channel

A previous study demonstrated the inhibitory effect of natural K^+^-conducting ChR (KCR1) on action potential firing *in vitro*^25^. We hypothesized that such channels hold promise for applications like seizure suppression *in vivo*. Therefore, we aimed to engineer a more efficient and highly sensitive KCR for *in vivo* applications, especially for transcranial activation. We started by incorporating KCR1 with multiple plasma membrane-targeting cassettes including the cleavable N-terminal signal peptide Lucy-Rho (LR), the ER export signal (E), and the plasma membrane trafficking signal (T) together with eYFP (Y)(**Fig. 1a**), which were found to significantly enhance the photocurrents of other ChRs^26,27^, resulting in iKCR (improved KCR). Notably, the fusion with the membrane-targeting module significantly enhanced the expression level of KCR1 and increased its photocurrent 10-fold compared to wild type KCR1 (referred as KCR1-eYFP, **Extended Data Fig. 1a, b**).

**Fig 1.**
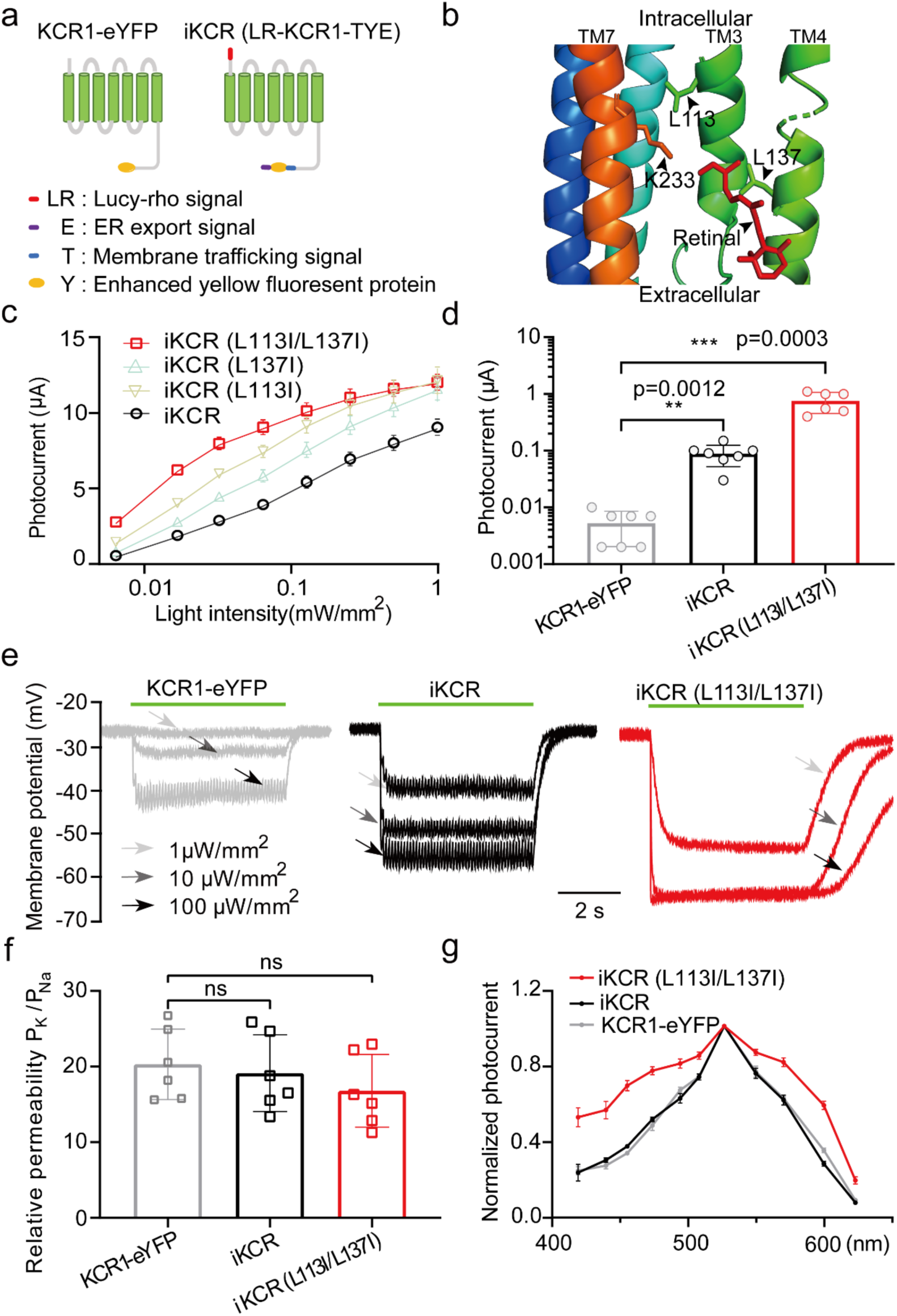
Engineering and characterization of light-gated K^+^ channels with high light-sensitivity in *Xenopus* oocytes. **a,** Schematic representation of KCR1-eYFP and iKCR constructs. **b,** The relative positions of amino acid residues L113, L137, and K233 in the KCR1 structure. K233 is covalently linked to all-trans retinal in the dark state. **c,** Light sensitivity of iKCR and its point mutants. Photocurrent amplitudes of KCR1 and optimized KCR1 variants measured by two-electrode voltage-clamp at a holding potential of ™40mV in Ringer solution (in mM: 110 NaCl, 5 KCl, 5 HEPES, 1 MgCl2, pH=7.6) with 2 mM BaCl2. The irradiation condition was 1 s, 530 nm green light. n=8-12 oocytes. **d,** Comparison of photocurrents among KCR1-eYFP, iKCR, and iKCR (L113I/L137I) upon illumination with dim green light (1 μW/mm^2^). **e,** Representative traces of membrane potential changes in KCR1-eYFP-, iKCR-, and iKCR (L113I/L137I)-expressing *Xenopus* oocytes upon 10 ms, 10 Hz 530 nm light stimulation with 1 μW/mm^2^ (light grey arrows), 10 μW/mm^2^ (grey arrows) and 100 μW/mm^2^ (dark grey arrows). **f,** Comparison of the relative potassium to sodium ion permeability, n=6 oocytes. **g,** Action spectra of KCR1-eYFP, iKCR, and iKCR (L113I/L137I), n=6 oocytes. Statistical analysis was performed by unpaired Mann Whitney test. Error bars are standard errors of the mean (SEM).

To further enhance its light sensitivity for transcranial optogenetic activation, we modified specific amino acids - L113 and L137 predicted to affect the gating and conductance of the channel. They are both located in the retinal binding pocket of KCR1, which might influence currents by modulating the photocycle **(Fig. 1b)**. We discovered that substituting Isoleucine (I) for Leucine (L) at either position 113 (relative to ChR2 I131^28^) or 137 (relative to ChR2 D156^29,30^) increased light sensitivity and photocurrents relative to iKCR and other mutations tested (**Fig. 1c, d** and **Extended Data Fig. 1c, d**). Interestingly, combining these mutations dramatically improved light sensitivity, generating around nine-fold larger photocurrents than iKCR and ninety-fold larger photocurrents than KCR1-eYFP under a dim irradiation condition (1 µW/mm^2^, **Fig. 1d**). In line with the larger photocurrents, iKCR (L113I/L137I) was more effective at hyperpolarizing oocytes than either iKCR or wildtype KCR1, reaching the reversal potential already with 10 µW/mm^2^ vs. 100 µW/mm^2^ for iKCR (**Fig. 1e**). Of note, the L113I and L137I mutations did not significantly affect the K^+^ selectivity of KCR1 (**Fig. 1f** and **Extended Data Fig.1e-h)**. Besides, iKCR (L113I/L137I) kept similar action spectrum peak as KCR1 and iKCR, but gave broader range of action spectrum (**Fig. 1g**). Given long wavelength light passes deeper into tissue, higher sensitivity at red spectrum for iKCR (L113I/L137I) may enable it to hyperpolarize cells located in the deep tissue. In summary, iKCR (L113I/L137I) exhibited much improved performance with better expression and higher light sensitivity in oocytes, potentially allowing its use for noninvasive inhibition *in vivo*. We name this channel as hsKCR for <u>h</u>ighly <u>s</u>ensitive <u>K</u>^+^ <u>c</u>hannel<u>r</u>hodopsin.

We next characterized hsKCR in mammalian neurons. hsKCR expression was confined to the soma and dendrites of rat organotypic hippocampal CA3 neurons (**Fig. 2a**). Without photoactivation, basic electrophysiological properties of the neurons transfected by either hsKCR or iKCR were unchanged comparing to nontransduced cells at the resting state in terms of series resistance (Rs), membrane resistance (Rm), holding current (Ihold) when clamped at ™74 mV, resting membrane potential (RMP) and action potential firing threshold (AP thresh.) (**Extended Data Fig. 2a**). Outward photocurrents were observed when hsKCR-expressing neurons clamped at ™60 mV were stimulated by the light with its wavelength spanning from 470 nm to 630 nm (**Fig. 2b**), suggesting hsKCR can be functionally photoactivated in neurons. Moreover, short green light pulses effectively inhibited action potential firing of the CA3 pyramidal neurons expressing hsKCR evoked by 1 s current injection (**Fig. 2c)**. This effect was more pronounced when the light intensity was more than 0.1 mW/mm^2^. The neuronal firing promptly recovered after photoactivation (**Fig. 2d**). Similar inhibitory effects were observed in iKCR-expressing CA3 pyramidal neurons but with 10 times more DNA electroporated comparing with hsKCR DNA transduced (10 ng/µl vs. 1 ng/µl at the same volume, **Extended Data Fig. 2b-d**).

**Fig 2.**
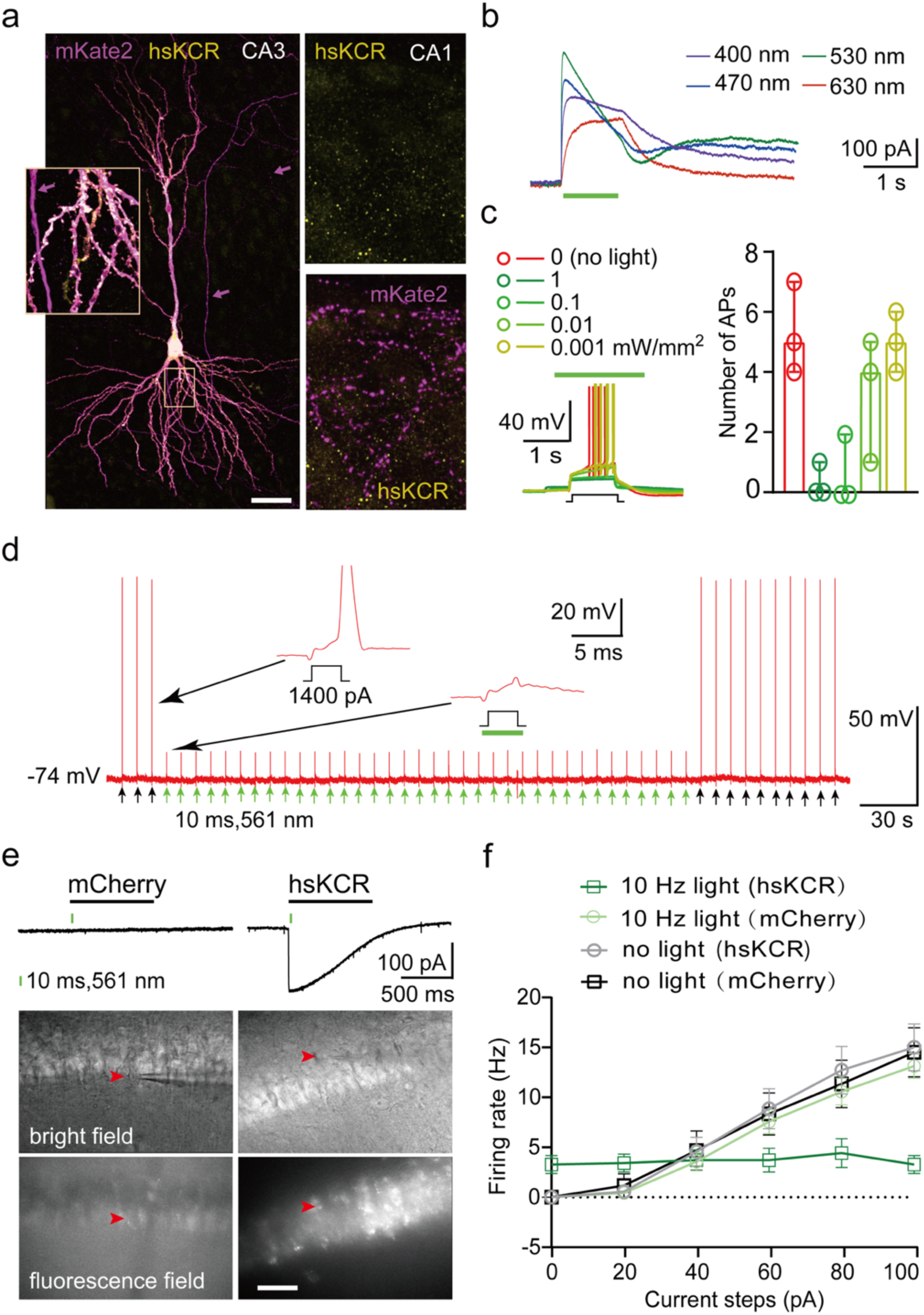
Photoactivation of hsKCR effectively silenced neuronal firing in rat hippocampal slice cultures (a-d) and mouse acute hippocampal slices (e-f). **a,** Confocal images of a CA3 neuron expressing hsKCR (yellow, anti-YFP staining) and mKate2 (magenta, anti-mKate2 staining). Note the absence of hsKCR immunostaining in axons (magenta arrows) in CA3 or in projections to CA1 (right panels). The same scale bar indicated 50 µm for the overview and 10 µm for the insets. **b,** Representative photocurrents recorded at ™60 mV in an hsKCR-expressing CA3 pyramidal neuron activated by the light with different wavelengths. The green bar indicates 1 s, 1 mW/mm^2^ light illumination. **c,** Exemplary current-clamp recordings of action potentials from an hsKCR-expressing CA3 neuron during 1 s current injection (black step, left panel). The green bar indicates 1 s application of 530 nm light at various intensities. Statistical analysis of the relationship between the number of action potentials and light intensities (right panel, median and interquartile range, p=0.003, Friedmann test, n=3 cells). **d,** Action potentials of an hsKCR- expressing neuron. Every 5 s, 3 ms-current steps (1400 pA) were injected to evoke single action potentials (arrows below traces, black steps in inserts). Starting from the 4^th^ pulse, green light flashes were triggered 1 ms before the current steps (green arrows/green bar, 5 ms, 530 nm, 10 mW/mm^2^). **e,** Representative photocurrents and fluorescence images of CA1 pyramidal neurons infected by mCherry (control) or hsKCR-mCherry viruses in whole-cell patch clamp experiments. Image scale bar: 100 μm. **f,** Quantification of firing rates in CA1 pyramidal neurons expressing hsKCR-mCherry or mCherry with or without 561 nm light illumination. (n=6-8 neurons from 3 mice). An optical fiber conducting 561 nm laser light was guided to the clamped cells for illumination.

Given that hsKCR showed higher red-shifted activation, larger photocurrents, and increased light sensitivity comparing with iKCR, we selected hsKCR for further investigation. hsKCR specifically expressed in mouse CA1 or CA3 pyramidal neurons two weeks after infection by AAV2/9-CaMKIIα-hsKCR-mCherry (**Extended Data Fig. 2e**). Photocurrent was detected upon green light simululation indicating its functional expression in the hippocampus (**Extended Data Fig. 2f**), without altering the reversal potential or membrane resistance of the infected neurons (**Extended Data Fig. 2g**). We injected current to these cells to evoke action potentials and tested if hsKCR photoactivation can inhibit the neuronal firing. As expected, we observed a significant light-induced reduction of the firing frequency in both hippocampal CA1 and CA3 neurons expressing hsKCR, while no such effect was observed in the brain slices infected by the control virus - AAV2/9-CaMKIIα-mCherry (**Fig. 2e, f** and **Extended Data Fig. 2h, I**). Thus, hsKCR can be functionally expressed in hippocampal neurons *in vivo*, and its activation effectively silenced neuronal firing.

### Anti-seizure effects of hsKCR

To assess the performance of hsKCR *in vivo*, we applied it in the context of epileptic seizures in mice. We expressed hsKCR using AAV2/9-CaMKIIα-hsKCR-mCherry virus in the mouse hippocampi and investigated whether optogenetic activation of hsKCR can inhibit hyperactive firing to alleviate epileptic seizures. We implanted an opto-tetrode bundle above the injection site for optical illumination and recording local field potential (LFP) (**Fig. 3a**). Alternatively, AAV virus encoding genetical Ca^2+^ indicator GCaMP7f was also injected to the same brain regions with a light guide implanted for in vivo Ca^2+^ imaging with fiber photometry. Additionally, a cannula was implanted contralaterally at the dentate gyrus for KA injection to induce seizures. We unilaterally injected KA into the dorsal hippocampi of mice to generate an acute pre-clinical seizure model^31-33^ (**Fig. 3a**). The induction of epilepsy was confirmed by the detection of the neuronal activation marker c-Fos and recordings of Ca^2+^ signals or LFPs *in vivo^34^*. Strong c-Fos expression was observed throughout the hippocampal formation in mice treated with KA but not the saline-treated ones, indicating increased neuronal activity in response to KA administration (**Extended Data Fig. 3a, b**). Behaviorally, KA-treated mice exhibited convulsive seizures (**Extended Data Movie 1**), and all mice studied displayed seizure-associated large Ca^2+^ signals (**Extended Data Fig. 3c**), either within focal seizures (FS, seizure scores 1–3) or generalized seizures (GS, seizure scores >3). Furthermore, LFP recordings using tetrodes also demonstrated seizures corresponding to behavioral seizure observations after KA administration (**Extended Data Fig. 3d**). These results confirm that the KA-induced seizure model was successfully established in mice.

**Fig 3.**
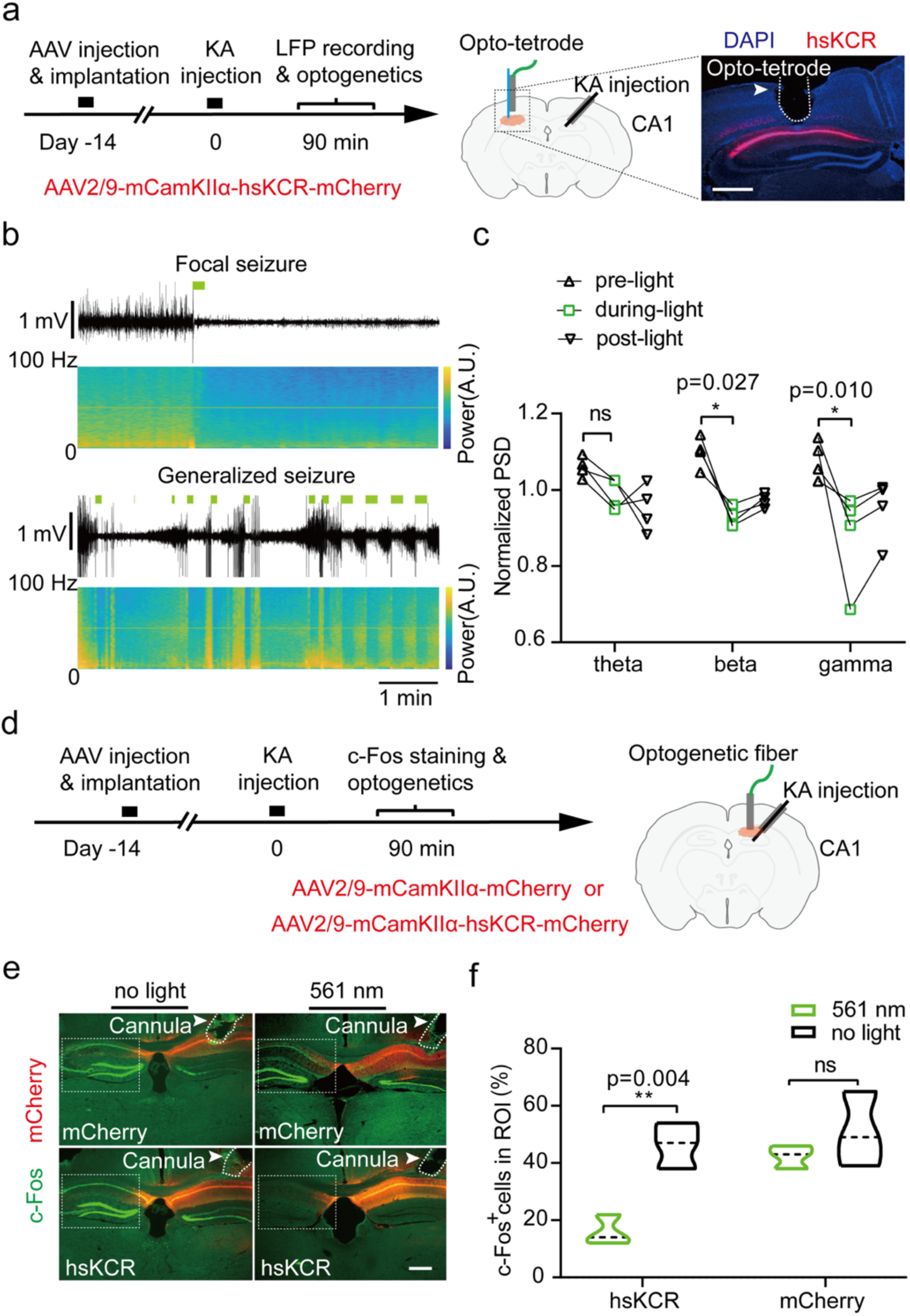
hsKCR-mediated suppression of KA-induced hippocampal seizure activity. a, Experimental schema of LFP recordings and optogenetics. Left panel: experiment time course. Right panel: a cartoon coronal section illustrates the arrangement of virus injection and cannula or Opto-tetrode implantation. An Opto-tetrode was implanted over the dorsal CA1 at the site of AAV2/9-mCaMKIIα-hsKCR-mCherry virus injection for optogenetic modulation and LFP recording, and a cannula was implanted into the contralateral dentate gyrus of mice for KA delivery. The mCherry immunostaining image verified hsKCR-mCherry expression just below the site of the opto-tetrode. Scale bar: 200 μm. **b,** Representative LFP traces during KA-induced FS and GS and corresponding power spectral analysis in hsKCR- expressing mice. Green bars indicate 10 Hz green light illumination. **c,** Normalization of mean PSD before, during, and after 10 Hz light illumination. Multiple paired *t*-test, n=4 mice. **d,** Experimental design of optogenetics and c-Fos immunofluorescence. An optogenetic fiber was implanted over the dorsal CA1 at the injection site of mCherry or hsKCR viruses, and a cannula was implanted into the ipsilateral dentate gyrus of mice for KA injection. **e** and **f,** Representative images (**e**) and quantifications (**f**) of c-Fos expression levels in hsKCR- and mCherry-expressing mice with or without green light illumination during 1.5 h of KA kindling. Two-way ANOVA followed by Sidak’s multiple comparison test was conducted. n=3 mice, scale bar: 200 μm.

Next, we tested whether optogenetic inhibition with hsKCR could decrease seizure activity using the above experiment settings. We recorded LFPs and monitored behavior for an initial 15-min baseline period, followed by KA injection. Subsequently, intermittent 10Hz illumination with 561nm light pulses was applied. Light transiently decreased both the magnitude and frequency of LFPs during seizure kindling (**Fig. 3b** and **Extended Data Fig. 3e**). Quantification of power spectral density (PSD) revealed a significant decrease in beta to gamma power during 10 Hz illumination (**Fig. 3c**). Furthermore, green light activation of hsKCR reduced hippocampal c-Fos expression 1.5 h after KA kindling (**Fig. 3d-f**). In contrast, KA-induced c-Fos expression did not change in the mCherry-expressing mice illuminated by the green light or the hsKCR-expressing mice without light stimulation, suggesting the inhibitory effects were specifically conducted by hsKCR photoactivation. Thus, hsKCR activation effectively inhibits KA-induced epileptic activity in mice.

We further assessed whether the anti-seizure effect of hsKCR expressed in the hippocampal CA1 may affect neuronal Ca^2+^ transients and seizures at the behavioral level. To this end we performed video recordings of the mice while simultaneously monitoring their Ca^2+^ activity on the side contralateral to KA injection in GCaMP7f-expressing CA3 pyramidal neurons using fiber photometry (**Fig. 4a and Extended Data Fig. 4a**). For paired comparison, seizures were induced twice by KA administration with 3 days interval in the mice expressing hsKCR or mCherry using a cross-over design. During seizure induction, mice were either treated first with or first without 561-nm green light illumination. In an additional group of mice, seizures were induced only once and the mice were randomly assigned to either a light-only or no light group for comparison.

**Fig 4.**
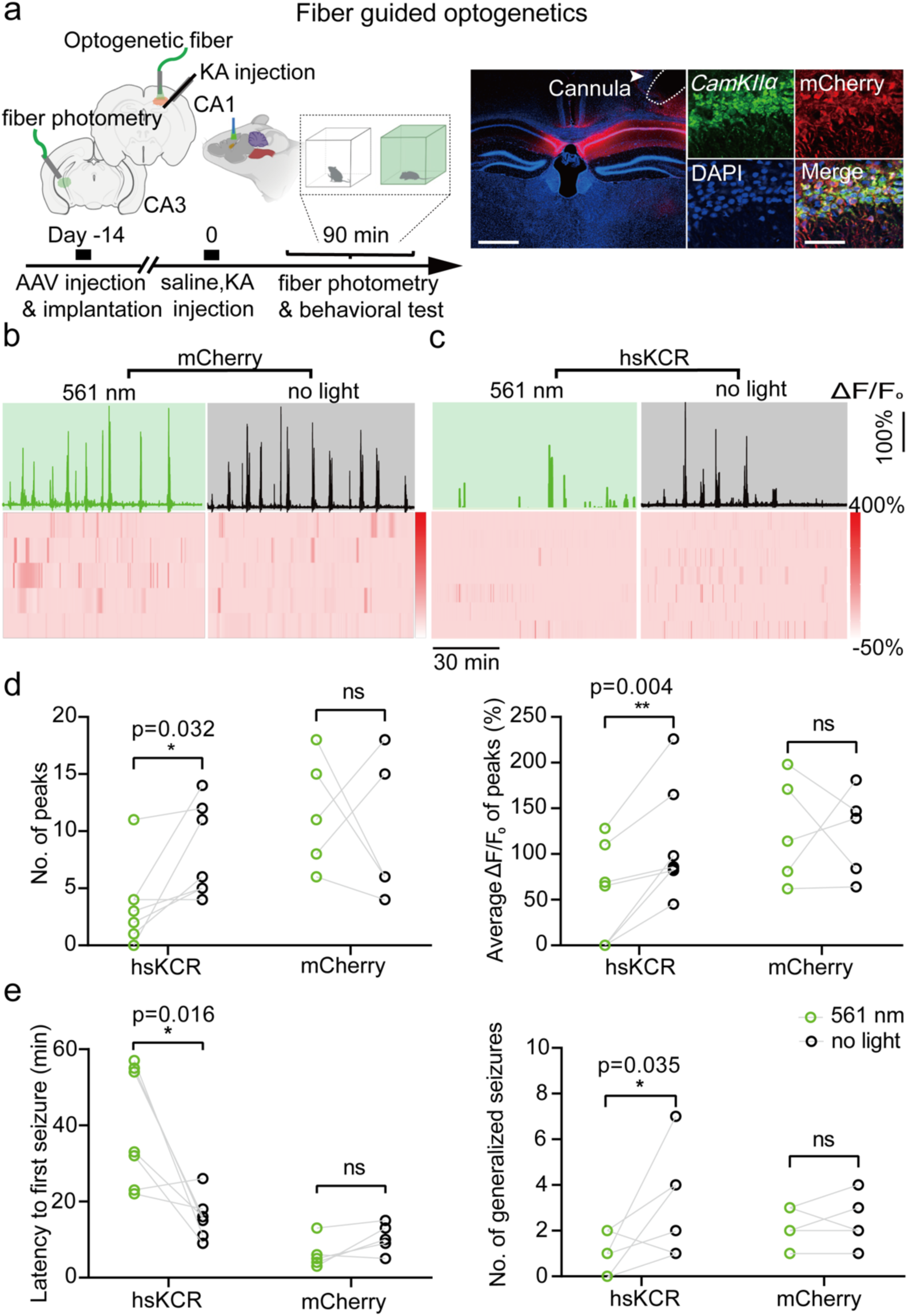
hsKCR effectively suppressed KA-induced convulsive seizures. a, Left: schema displaying our fiber-guided optogenetic intervention to modulate KA-induced acute seizures. AAV2/9-mCaMKIIα-hsKCR-mCherry or AAV2/9- mCaMKIIα-mCherry viruses were injected into the CA1 of one hemisphere, while AAV2/9-mCaMKIIα-GCaMP7f was injected into the CA3 of the other hemisphere. Right: expression of hsKCR in CA1 neurons, visualized by immunofluorescence against mCherry. Scale bars: 200 μm in the overview and 100 μm in the inset. **b** and **c,** Top: representative Ca^2+^ activity from **(b)** AAV2/9-mCaMKIIα-mCherry-expressing or **(c)** AAV2/9-mCaMKIIα-hsKCR- mCherry-expressing mice with (left) or without (right) 561 nm green light illumination during a 1.5 h period after KA kindling. Bottom: heatmaps of Ca^2+^ activities in CA3 regions. Each row represents the Ca^2+^ signal of an individual mouse, with a total of five **(b)** or seven **(c)** mice shown. The color scale indicates ΔF/F0, with warmer colors indicating higher fluorescence signals. **d,** Quantification of seizure activity indicated by the average peak Ca^2+^ signals (ΔF/F0 > 40%) and the number of peaks in these mice. **e,** Quantification of behavioral seizures is indicated by the latency to the seizure onset and the number of generalized seizures in these mice. Illumination condition: 561 nm laser, 10 ms light pulse at 10 Hz with a cyclic light pattern of 10 s-on/10 s-off. The light power was 6 mW (180 mW/mm^2^) measured at the fiber tip. Statistical analysis was performed using paired *t*-test, and error bars represent the SEM.

In Ca^2+^ recording experiment, we observed similar anti-seizure effects of hsKCR as the LFP measurement. In the saline injection group, the baseline of Ca^2+^ activity (refer to ΔF/F0 > 10%) did not show obvious changes upon photoactivation in both hsKCR- and mCherry-expressing mice (**Extended Data Fig. 4b**). It suggests that hsKCR activation has no apparent influence on Ca^2+^ activity at the basal condition. In the KA group, Ca^2+^ signals were upregulated as expected due to seizure induction. Upon 561nm light illumination, such signals were efficiently suppressed in hsKCR-expressing mice, but it was not observed in the mCherry mice (**Fig.4c**). Subsequently, we analyzed Ca^2+^ peaks with ΔF/F0 more than 40%, which were only observed in KA-treated mice thus highly associated with seizure responses. In mice expressing hsKCR, green light illumination reduced the Ca^2+^ peak numbers by more than half compared to the no-light condition (green light: 3.57±1.36 vs. no light: 8.14±1.53; **Fig. 4d**). Additionally, the average ΔF/F0 change of the Ca^2+^ peaks was also significantly decreased in the presence of green light (green light: 51.3±20.5 vs. no light: 112.3±23.3; **Fig. 4d**). On the contrary, in the mCherry-expressing mice green light illumination induced no significant difference of either the number of Ca^2+^ peaks (green light: 11.6±2.20 vs. no light: 9.82±2.80; **Fig. 4d**) or the average ΔF/F0 (green light: 125.2±26.0 vs. no light: 123.0±21.4; **Fig. 4d**). Thus, optogenetic activation of hsKCR suppressed hyperactive Ca^2+^ signals during seizures without apparently affecting basal Ca^2+^ signals indicating that it can inhibit neural hyperactivity.

Regarding the seizure behavior, close observation revealed that green light irradiation significantly prolonged the latency to seizure onset in hsKCR-expressing mice of the paired study, from 15.9±2.06 min to 39.4±5.84 min (**Fig. 4e**). In contrast, light did not delay seizure onset in mCherry-expressing mice (green light: 6.20±1.72 min vs. no light: 9.02±1.72 min, **Fig. 4e**). Notably, the inhibitory effect of hsKCR photoactivation caused a pronounced decrease in the number of GS by the behavior observation (**Fig.4e**). While on average hsKCR-expressing mice with light stimulation had less than one GS bout (0.86±0.34), those that did not receive light exhibited more than three GS bouts (3.29±0.81). As expected, illumination had no obviously modulatory effect on seizure activity in the mCherry-expressing mice. We also observed similar inhibitory effect on seizure behavior in the unpaired experiment, in which both seizure onset and bouts of FS and GS were suppressed upon light illumination in the hsKCR-expressing mouse group (**Extended Data Fig. 4c**). Overall, these findings strongly suggest that photoactivation of hsKCR effectively inhibits seizure behavior in our acute seizure model.

### Suppressing seizures with bilateral transcranial optogenetic stimulation of hsKCR

The large photocurrents, high light sensitivity, and seizure-suppressing properties of hsKCR suggested it may be an ideal candidate for deep transcranial optogenetic inhibition, which holds great potential for noninvasive applications in clinic^35^. Transcranial photoactivation of hsKCR can minimize the tissue damage to a large extent since the optical fiber is not implanted into the brain anymore. However, more light power may be required due to the power decay in the light path through the brain tissue, which in turn may heat up the local brain as a side-effect^7^. We indeed detected such temperature increase caused by the optogenetic light source in our experiment settings **(Extended Data Fig. 5a**). Thus, we compared the safety aspects of intracranial and transcranial photoactivation, including inflammation and tissue heating effects (**Fig. 5a**). Notably, around the implantation site of the optical fiber, we observed a significant increase of Iba-1 and Gfap immunoreactivity (**Fig. 5b** and **Extended Data Fig. 5b, c**). It suggests there was activation of microglia and astrocytes adjacent to the injury site, indicating a pronounced inflammatory response due to the local tissue damage^36^. Iba-1 and Gfap staining kept at a low level in the brains of mice that did not undergo fiber implantation surgery. Even transcranially illuminating at a high light power, there was no remarkable increase in Iba-1 or Gfap expression (**Fig. 5b** and **Extended Data Fig. 5b, c**). Besides, regarding the heating effect, light stimulation using implanted optical fibers caused a slight increase at both CA1 and skull surface, which did not occur in CA1 area during transcranial illumination (**Fig. 5c** and **Extended Data Fig. 5d**). Thus, transcranial optical stimulation minimizes the risk of local brain inflammation and tissue heating effects.

**Fig 5.**
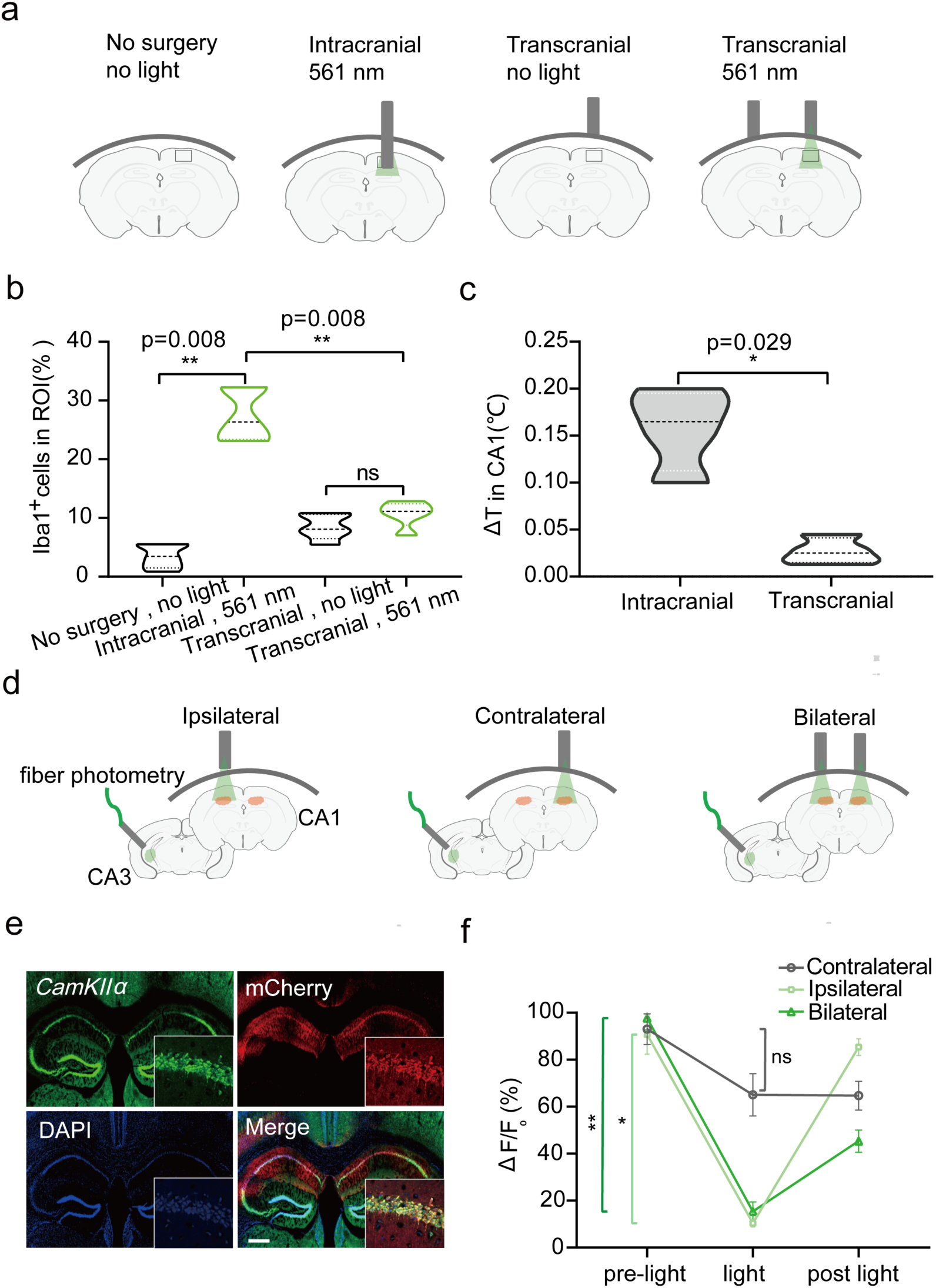
hsKCR-conducted noninvasive deep brain optogenetic inhibition while minimizing tissue injury. **a,** Experimental schema to evaluate the safety of intracranial and transcranial optogenetic stimulation. **b,** Quantification of the percent of Iba^+^ microglia among all cells (DAPI^+^) in the regions indicated by the black boxes shown in **(a)**. The stimulation protocol involved a cyclic light pattern: 50 ms 561 nm light pulses at 10 Hz for 10 s followed by 10 s without illumination. Statistical analysis was performed using one-way ANOVA with Tukey’s multiple comparisons test, n=5 mice per group. **c,** Temperature changes during intracranial and transcranial optogenetic stimulation in the CA1. A comparison between the two approaches was made using unpaired Mann-Whitney test, n=4 mice per group. **d,** Schematic illustration of contralateral, ipsilateral, and bilateral transcranial optogenetic stimulation during pilocarpine-induced seizures. **e,** Fluorescent images showing the coexpression of mCherry (hsKCR indicator) and CaMKIIα (a marker for pyramidal neurons) in a brain slice after bilateral injection of hsKCR AAVs. Scale bar, 100 μm. **f,** Quantification of normalized Ca^2+^ signal changes before, during, and after different ways of light illumination. The Ca^2+^ change was indicated as ΔF/F0 (%). Ca^2+^ signals correlated with first GS behaviors induced by pilocarpine as indicated by the arrows in red.

After safety evaluation of our noninvasive optical stimulation protocol, we characterized the inhibitory effect with transcranial illumination on different acute seizure models. We first examined whether activation of hsKCR by transcranial illumination could inhibit neural activity in pilocarpine-induced persistent seizure, which is a well-established model of clinical status epilepticus^37^. We bilaterally injected AAV2/9-mCaMKIIα-hsKCR-mCherry virus into the CA1 region to transduce excitatory hippocampal neurons. To monitor seizure activity in real-time, we also injected AAV2/9-mCaMKIIα-GCaMP7f virus into the hippocampal CA3 and detected Ca^2+^ increase in pilocarpine-induced persistent seizure (**Extended Data Fig. 5e**). In the meantime, we applied transcranial green light illumination to the contralateral, ipsilateral, and bilateral sides of the hippocampus relative to the Ca^2+^ recording site (**Fig. 5d**). Histological staining indicated that hsKCR was expressed in the hippocampal pyramidal neurons (**Fig. 5e**). Interestingly, we observed significant Ca^2+^ decrease during ipsilateral optogenetic transcranial intervention (**Fig. 5f**). In comparison, contralateral photoactivation of hsKCR only induced mild Ca^2+^ decrease. It suggests that local light stimulus of hsKCR might cause more remarkable inhibition than contralateral illumination. To check if there may be some synergetic effect by illumination on both sides of the hippocampus. We compared the inhibitory effect of bilateral illumination with ipsilateral stimulus. Bilateral optical stimulation potentiated such effect by delaying the recovery of Ca^2+^ signal after illumination, but did not induce more pronounced Ca^2+^ decrease during the light stimulus (**Fig. 5f**). Given epileptic activity rapidly propagates to form a widely distributed neural network^38^, the delayed recovery effect may result from such network connection. In summary, bilaterally transcranial activation of hsKCR caused robust reduction of epileptic Ca^2+^ activity.

Further, we took the KA-induced epilepsy model to assess if <u>b</u>ilateral <u>t</u>ranscranial <u>o</u>ptogenetics (BTO) may alleviate seizures at the behavior level. After hsKCR was bilaterally expressed in CA1 hippocampal neurons via AAV injection, a guide cannula was inserted into one side of the hippocampal CA1 region for KA delivery **(Fig. 6a)**. The cannula was embedded at a 30-degree angle to not block the transcranial light path with light guides bilaterally mounted on top of the mouse skull. Besides, AAV2/9-mCaMKIIα-GCaMP7f virus was injected into the hippocampal CA3 region with a light guide implanted for Ca^2+^ recording using fiber photometry. After KA administration, we observed that spontaneous seizures were effectively inhibited by transcranial green light in hsKCR-expressing mice (**Extended Data Fig. 6a, b**). The latency to the first seizure onset was significantly delayed by green light stimulation (22.4±5.04 min) compared to no light (10.4±1.56 min; **Fig. 6b**). Additionally, the number of generalized seizure bouts was significantly reduced (0.71±0.36) compared to no light condition (2.49±0.57; **Fig. 6c**). We also observed similar inhibitory effect on the total number of seizure bouts (**Extended Data Fig. 6c**). In contrast, in mice injected with AAV2/9- mCaMKIIα-mCherry control virus, green light stimulation had no effect on the latency to the first seizure (green light: 11.8±2.44 min vs. no light: 14.4±2.42 min) or generalized seizure bouts (green light: 1.80±0.58 vs. no light: 2.42±0.24) as well as the total seizure numbers (**Fig. 6b, c** and **Extended Data Fig. 6c**). Therefore, in accordance with the BTO inhibitory effect on epileptic Ca^2+^ signals, transcranial photoactivation of hsKCR also efficiently inhibits KA-induced seizure onset in mice.

**Fig 6.**
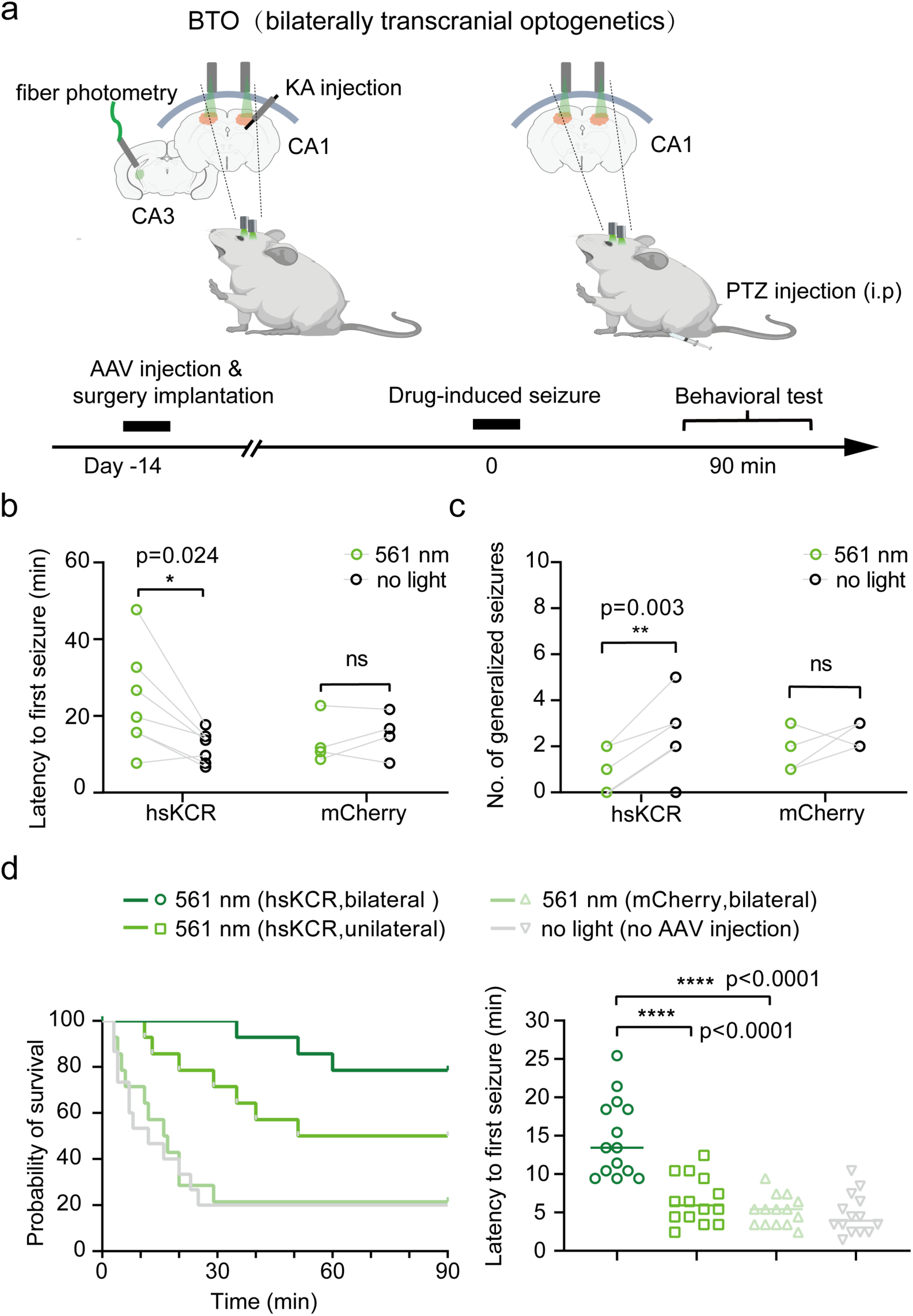
*In vivo* anti-seizure effects of bilateral transcranial optogenetics in different seizure models. a, Schema depicting our BTO approach for controlling KA and PTZ-induced seizures. **b,** Quantification of behavioral seizures indicated by the latency to the first seizure during KA kindling. mCherry mice n=4, hsKCR mice n=7. Statistical analyses were performed using multiple paired *t*-test. **c,** Quantification of behavioral seizures indicated by the numbers of GS during KA kindling in the same experiment as (b). **d,** Analysis of the survival rate and latency to first seizure induced by PTZ treatment. The transcranial stimulation protocol involved a cyclic light pattern that delivered 50 ms light pulses at 10 Hz for 10 s followed by 10 s gap without illumination. The power of the applied 561 nm light was approximately 15 mW (measured at the outlet tip of the optical fiber). Statistical analysis was performed using one-way ANOVA with Tukey’s multiple comparisons test. Survival rates were analyzed with a Kaplan-Meier curve with the log-rank (Mantel-Cox) test. n=14 mice per group.

Finally, to further validate the antiepileptic effect of BTO in combination with hsKCR, we utilized the PTZ-induced seizure model^39^. PTZ-induced epilepsy is highly lethal if seizure activity is not promptly inhibited^40,41^. We recorded PTZ-induced seizure behavior and analyze the survival rate. As expected, mice expressing the control virus that underwent bilateral transcranial illumination exhibited a similarly poor survival rate as non-transduced mice after PTZ injection (**Fig. 6d**). But interestingly, hsKCR-expressing mice demonstrated a significant improvement in survival rate and prolonged latency to the first seizure compared to control mice under the same BTO conditions (**Fig. 6d**). Unilateral transcranial illumination of hsKCR-expressing mice also meliorated PTZ-induced seizure mortality, but the survival rate was still much lower than the BTO mouse group, and the time to first seizure was no longer prolonged. BTO using hsKCR also reduced the Racine scores to the greatest extent **(Extended Data Fig. 6d**). Therefore, the large photocurrent, highly light-sensitive potassium-selective channelrhodopsin hsKCR can be sufficiently activated with bilateral transcranial illumination to inhibit seizure activity in three different mouse models of epilepsy.

## Discussion

We developed a potent inhibitory optogenetic tool – hsKCR with great light sensitivity, and showed its activation could efficiently suppress neural activity in brain slices and KA-induced seizure model of mice. Such inhibitory effect could be maintained by transcranial activation of hsKCR in three different seizure models. Especially, BTO intervention of hsKCR in the hippocampus resulted in dramatic increase of mouse survival rate of PTZ-induced epileptic seizure. Thus, we introduced a novel transcranial optogenetic therapy for the treatment of epilepsy potentially. To the best of our knowledge, this study is the first to demonstrate the effectiveness of transcranial optogenetic inhibition in controlling epilepsy by modulating potassium ion efflux in awake mice. This approach offers precise temporal control over neural spiking, allowing targeted inhibition of abnormal neuronal activity. Furthermore, our method eliminates the need for invasive procedures such as implanting light-emitting devices into brain tissue^42^, thereby reducing potential risks and complications. In addition, the ultra-high light sensitivity of hsKCR requires much less light power, thus minimizing the side effects of local tissue heating.

Derived from the natural light-gated KCR1^25^, its engineered variant hsKCR demonstrated larger photocurrents and higher light sensitivity, while maintaining selectivity for potassium ions. Together with the enhanced light sensitivity, the closing kinetics of hsKCR was prolonged from 48 ms to 824 ms (**Extended Data Fig. 1d**), which likely contributes to its efficacy in suppressing seizures. In contrast, a previous study reported that PAC-K, the combination of a blue light-activated adenylyl cyclase bPAC and a cAMP-dependent potassium channel SthK, exacerbated rather than prevented epilepsy^43^. An additional light-activated K^+^ channel WiChR was recently discovered^44^. As WiChR has a higher selectivity for K^+^ ions than KCR1, it will also be an attractive candidate for future testing in epilepsy models.

Long-term pharmacological treatment for epilepsy often faces limitations in terms of efficacy, leading to drug resistance^45,46^. Surgical resection is only feasible when the epileptogenic zone is sufficiently far from critical brain regions^47^. Therefore, genetic therapies have emerged as a promising option for the treatment of epilepsy due to their ability to modulate neuronal excitability with greater precision than pharmacological tools^48,49^. Previous studies have demonstrated the effectiveness of overexpressing engineered K^+^ channels in reducing seizure frequency and duration in both focal neocortical and temporal lobe epilepsy models^50^. Compared to optogenetic approaches, such kind of methods offer the advantage of no requirement of additional optical stimulation. However, it lacks spatiotemporal precision and has limited reversibility^51,52^. Consequently, there are potential risks of depression if the neural activity balance is not precisely modulated with these methods. In our study, we present a novel optogenetic intervention for seizure control through bilateral transcranial optical activation of hsKCR. Its activation and deactivation can be well controlled by external light source with millisecond precision, providing a powerful tool for dynamically modulating neural activities in a reversible manner. Moreover, hsKCR might be sufficiently activated by red-light above 600 nm for BTO inhibition. Red light penetrates deeper tissue and causes less photo-induced damage effect than blue-green light. Additionally, the possibility of using BTO within a closed-loop neuromodulation strategy holds great promise for seizure control *in vivo*.

Translational application of optogenetics to deep brain regions faces many additional challenges when compared with more accessible organs like the retina, which in recent years has seen translational and clinical advances. Safe and effective gene targeting and light delivery need further development before optogenetics become a treatment option in clinic^35^. Recent advancements in engineering wireless and biocompatible μ-ILEDs, and upconversion of near-infrared nanoparticles may be promising solutions for deep brain optogenetic stimulation^53^. For example, to target the superficial cortical and subcortical brain regions, it is possible to deliver light from outside the meninges by miniaturized and battery-free light-emitting materials^42^. Moreover, novel noninvasive gene delivery technologies targeting the central nervous system have also emerged as viable options^54,55^. In light of these developments, hsKCR-mediated transcranial optogenetic inhibition holds significant promise for treating neurological and psychiatric disorders characterized by focal and pathological hyperactivity.

## Materials and Methods

### Animals

C57BL/6J mice, 8-10 weeks old, were obtained from Shanghai Model Organisms Center and raised at the animal facility of Southern University of Science and Technology. The mice were housed in a controlled environment with a 12-hour light/dark cycle, maintained at a temperature of 20-26°C and humidity of 30- 60%. They had access to food and water *ad libitum*. All experimental procedures were conducted in accordance with the guidelines of institutional animal care and use committee at Southern University of Science and Technology, following the National Institutes of Health guidelines for the care and use of laboratory animals.

Wistar rats (Envigo HsdCpb:Wu strain) were bred at the University Medical Center Hamburg-Eppendorf animal facility and sacrificed according to German Law (Tierschutzgesetz der Bundesrepublik Deutschland, TierSchG) with approval from the Behörde für Justiz und Verbraucherschutz (BJV)-Lebensmittelsicherheit und Veterinärwesen, Hamburg and the animal care committee of the UKE.

### Expression plasmids

The KCR1 gene was optimized for mouse expression and synthesized using GeneArt Strings DNA Fragments (Thermo Fisher Scientific) according to the published DNA sequence^25^. The synthesized KCR1 fragment was inserted into the pGEMHE vector with eYFP and plasma membrane-targeting cassettes including LR, E, and T to make the iKCR expression plasmid, using BglII and XhoI restriction sites. Or, the KCR1 fragment was inserted into the pGEMHE vector that contains the eYFP cassette to make the KCR1-eYFP plasmid, using BamHI and XhoI restriction sites. Point mutations were introduced using the QuikChange site-directed mutagenesis method to make hsKCR plasmid modified based on iKCR plasmid. The DNA sequence was confirmed by commercial Sanger sequencing service. The generated KCR1 variants, iKCR and hsKCR, were transferred to the AAV vectors (pAAV-CaMKIIα) using the compatible BamHI and HindIII restriction sites at the N- and C-terminus of the complete insert.

### Complementary RNA preparation for expression in *Xenopus* oocytes

After confirming the sequence through DNA sequencing, the pGEMHE plasmids carrying different KCR1 variants were linearized by NheI digestion and used for *in vitro* transcription of complementary RNA (cRNA) using the AmpliCap-Max T7 high-yield message maker kit (Epicentre Biotechnologies). For all the KCR1 expression variants, 30 ng of cRNA were injected into *Xenopus* oocytes. The cRNA-injected oocytes were cultured in ND96 solution containing 96 mM NaCl, 5 mM KCl, 1 mM MgCl2, 1 mM CaCl2, 5 mM HEPES, pH 7.4, and supplemented with 10 µM all-trans-retinal at 16°C. The laparotomy procedure to obtain oocytes from *Xenopus* laevis was conducted following the principles of the Basel Declaration and recommendations of Landratsamt Wuerzburg, Veterinaeramt. The protocol, approved by the responsible veterinarian, was carried out under license #70/14 from Landratsamt Wuerzburg, Veterinaeramt. Oocytes were imaged using a Leica DMi8 inverted microscope and a Leica DFC3000G CCD camera. For checking the eYFP expression in transfected oocytes, the excitation light wavelength was 490-510 nm, and the acquisition of the emission channel was 520-550 nm. The imaging process was conducted using Leica Application Suite X software (v2.0.14332.0).

### Two-electrode voltage-clamp recording

Electrophysiological measurements were conducted at room temperature (20-23°C) using a two-electrode voltage-clamp amplifier (TURBO TEC-03X, npi electronic GmbH, Germany). The specific bath solutions for each electrophysiological recording were indicated in the figure legends. Electrode capillaries (Ф = 1.5 mm, Wall thickness 0.178 mm, Hilgenberg) were filled with 3 M KCl. The light source was a 530 nm LED with adjustable powers (Thorlabs Inc.) used for optogenetic illumination. Data acquisition was performed using a USB-6221 DAQ device (National Instruments) and WinWCP software (v5.5.3, Strathclyde University, UK).

### Electrophysiological recordings in hippocampal slice cultures

The procedure for preparing organotypic cultures was modified from Stoppini *et al*, using a growth media without antibiotics^56,57^. Hippocampal CA3 neurons were transduced using single-cell electroporation cultured at 6-14 days *in vitro* as described^58^. Recordings from neurons electroporated with iKCR (pAAV- CaMKIIα-LR-KCR1-TYE) at 10 ng/µl were performed 6-12 days later. Neurons electroporated with 1ng/µl hsKCR (pAAV-CaMKIIα-KCR1-2.0-L113I-L137I) were recorded 9-17 days after the transfection. pCI-syn-mKates (20 ng/µl) was co-transfected in all neurons. For recording, slice cultures were transferred to the stage of an upright microscope (BX61WI Olympus, Japan) that was perfused with 30-31 °C solution containing (in mM): 119 NaCl, 26.2 NaHCO3, 11 D-glucose, 4 MgCl2, 2.5 KCl, 1 NaH2PO4, 4 CaCl2, pH 7.4, 308 mOsm/kg, saturated with 95% O2 and 5% CO2. D-CPPene (1 µM), NBQX (10 µM) and picrotoxin (100 µM) were added to block synaptic activity (Hello Bio). The intracellular solution contained (in mM): 135 K^+^ gluconate, 10 HEPES, 4 MgCl2, 4 Na2-ATP, 0.4 Na-GTP, 10 Na2-phosphocreatine, 3 ascorbate, pH 7.22, 296 mOsm/kg. The liquid junction potential (−14.4 mV) was measured and corrected. Series resistance was less than 15 MΩ. Whole-cell patch-clamp recordings were made using an Axopatch 200B (Molecular Devices, USA), National Instruments A/D boards and *Ephus* software running in MATLAB (v2018b). Photostimulation was applied through a water immersion objective (Plan-Apochromat, 40x 1.0 numerical aperture, Zeiss) using an LED (Mightex Systems, Canada) coupled via a dual camera port with a multimode fiber (1.0 mm) and collimator (Thorlabs, USA). A power meter fitted with a silicon detector (Newport 1936R, 818-ST2) was used to calibrate the light intensity in the specimen plane. Pyramidal neurons in CA3 were clamped at ™60 mV or recorded in the current-clamp mode. Data were analyzed using MATLAB scripts.

### Electrophysiological recordings in acute brain slice

Male C57BL/6J mice were decapitated, and brains were immediately dissected and immersed in an ice-cold solution containing (in mM): 30 NaCl, 26 NaHCO3, 10 D-glucose, 4.5 KCl, 1.2 NaH2PO4, 1 MgCl2, 194 sucrose, and saturated with 95% O2 and 5% CO2. Coronal slices containing the hippocampus were cut at 350 μm thickness for electrophysiology recording using a vibratome (VT1120S, Leica Systems, Germany). Brain slices were rinsed twice in artificial cerebrospinal fluid (aCSF) containing (in mM): 124 NaCl, 26 NaHCO3, 10 D-glucose, 4.5 KCl, 1.2 NaH2PO4, 1 MgCl2, 2 CaCl2, 30 sucrose, and bubbled with 95% O2 and 5% CO2. Slices were gently moved to a brain slice keeper containing aCSF saturated with 95% O2 and 5% CO2. Incubation of slices was at 34°C for 30 min before transferring to room temperature for at least 1 h prior to recording. Slices were then individually transferred to a recording chamber (RC26G, Warner Instruments, USA) fixed to the stage of an upright microscope (BX51W, Olympus, Japan). During recording or imaging, slices were continuously perfused with aCSF saturated with 95% O2 and 5% CO2 at a flow rate of 3 ml/min. We chose mCherry^+^ cells for whole-cell patch-clamp recordings on acute hippocampal slices.

In order to determine the reversal potential of hsKCR, the mCherry^+^ cells were held at a range of membrane potentials, starting from ™70 mV and incrementally increased by 5 mV up to ™35 mV. Subsequently, the cell was exposed to a 10 ms pulse of 561 nm light, resulting in a gradual transition of the inward current to an outward current as the membrane potential increased. The reversal potential was determined when there is no current upon the 561 nm light pulse. To measure cell membrane resistance, cells received a ™10 mV pulse when holding at ™70 mV.

To test the optogenetic inhibitory effect when activating hsKCR on neurons, the initial membrane potential was held at ™70 mV. Then a series of currents (from 0 pA to 110 pA) was injected into the mCherry^+^ cells while the action potentials were recorded. For optogenetic manipulation, the cells received 500 ms constant 561 nm light illumination or the light at 10 Hz for 10 ms. The recording pipette was filled with (in mM): 128 potassium gluconate, 10 NaCl, 10 HEPES, 0.5 EGTA, 2 MgCl2, 4 Na2ATP, and 0.4 NaGTP. The acquisition frequency was 20 kHz. Neuron spiking traces were imported into the Fitmaster software (HEKA Elektronik, Germany) for analysis.

### Electrophysiological recordings *in vivo*

Adjustable opto-tetrode micro-drives were used for *in vivo* optogenetic simulation and electrophysiological recordings. The drive configuration principally followed Kubie’s design with modifications^59^. The bases were CNC-machined aluminum parts, and each had a customized electronic interface board (EIB) mounted on the side, to avoid interference with the optical fiber. Vertical adjustment for the micro-drives was enabled by two sets of supporting screws and nuts. Each drive was loaded with closely aligned one optical fiber (230 um, RWD Life Science) and 4 tetrodes. The tetrodes were twisted from HLM-coated, Cr20/Ni80 electrode wires (17.5 um, Stablohm 650, California Fine Wire). Wire-to-EIB connections were established by inserting wires and gold-plated pins into the EIB pads. The exposed wire middle piece and optical fiber were secured on the base using UV adhesive. Before the surgery, the end of the tetrode bundle was cut to approximately 500 μm protruding the optical fiber, and gold-plated to the resistance of 200-250 kΩ using the NanoZ system (White Matter LLC). To minimize non-neuronal electromagnetic interference during recording, the EIBs had reference pads connected to the aluminum base and a female connector, which was further connected to a skull screw via a stainless-steel wire during the surgery.

Following recovery from surgery, mice were screened and recorded head-fixed on a running wheel. The screening procedure was conducted daily to ascertain if tetrode tips were located near the CA1 region. Both single-unit activity, local field potentials and the reactions to short optical stimulation were recorded and used to identify how much the drive needed to be advanced.

To record intracranial electrical signals, the animal was connected to an Open Ephys acquisition board by the EIB via a headstage (Intan Technologies), where the signal recorded was also amplified. After amplification, signals were visualized and recorded by the Open Ephys GUI (v0.6.4). For the recording of LFPs, signals were bandpass filtered between 1-300 Hz.

To calculate LFP powers, the raw LFPs were filtered to eliminate the 10 Hz harmonics and divided into theta (5-10 Hz), beta (12-30 Hz) and gamma (30-80 Hz) bands. For each animal, a few segments of the entire recording were selected (ranging from 10 to 40 min) and classified into pre-light (within 5 seconds before stimulation), during light and post-light (within 5 seconds after stimulation). Filtered and classified LFP data were squared and averaged for statistical analysis.

### Fiber photometry

Fiber photometry recording was carried out using a commercial device (R810, RWD Life Science, China). The excitation wavelengths at 470 nm and 410 nm were used to detect the GCaMP7s fluorescence indicating intracellular Ca^2+^ changes and its isosbestic point for motion control, respectively. In brief, 470 and 410 nm laser beams were first launched into the fluorescence cube and then into the optical fibers. The fluorescence was collected by a CMOS camera at 200 Hz. *In vivo* recordings were carried out in an open-top mouse cage (L-W-H: 21.6 × 17.8 × 12.7 cm). The photometry signal F was derived as F470/F410, and Ca^2+^ changes were calculated as ΔF/F0 = (F – F0)/F0, where F0 is the median of the photometry signal within a defined time window. The average of peak ΔF/F0 values and the number of events for each mouse were analyzed.

### Animal surgeries and stereotaxic injections

Mice were anesthetized with sodium pentobarbital (100 mg/kg, i.p., Sigma-Aldrich), or with isoflurane gas (RWD Life Science, China) for both induction and maintenance of anesthesia throughout the surgery. Surgical procedures were conducted using a standard stereotaxic apparatus (RWD Life Science, China). The body temperature of mice was maintained using a thermostatic heat blanket (RWD Life Science, China), and mouse eyes were protected with moisturizing ointment. An incision was made to the mouse’s head to expose the skull surface and disinfected prior to the incision. Burr holes were stereotactically drilled on the skull.

For optogenetic manipulation, AAV2/9-mCaMKIIα-hsKCR-mCherry virus (5× 10^12^ vg/ml, 200 nl) was injected into the hippocampal CA1 region (AP −2.0, ML −1.3, DV −1.2 mm) in 8-week-old wildtype mice. AAV2/9-mCaMKIIα-mCherry virus (5× 10^12^ vg/ml, 200 nl) was injected in the contral mouse groups. All AAV viruses used in this study were purchased from Taitool Bioscience Co.Ltd. (Shanghai, China). An adaptor for a 200-μm, 0.39-NA optical fiber (RWD Life Science, China) was implanted above the injected site. In the transcranial optogenetic experiment, AAV2/9-mCaMKIIα-hsKCR-mCherry virus was bilaterally injected into hippocampal CA1 regions (AP −2.0, ML ±1.3, DV −1.2 mm) with injection route tilted by 30-degree angle, so that two optical fiber adaptors can be mounted to the skull surface above the injection sites with dental cement afterwards.

For volumetric Ca^2+^ recording using fiber photometry, AAV2/9-mCaMKIIα-GCaMP7f virus (5× 10^12^ vg/ml, 200 nl) was unilaterally injected into the hippocampal CA3 (AP −2.9, ML −3.2, DV −3.5 mm). An adaptor for a 200-μm, 0.39-NA optical fiber (RWD Life Science, China) was implanted above the injected site or tilted by 30-degree angle.

For *in vivo* LFP recording, a circular section of skin was removed to expose the skull. Once the skull surface was clean and flat, bregma and lambda were clear and aligned. Two skull screws (M1) were implanted respectively at the frontal bone and the interparietal bone. A cannula was implanted into the contralateral dentate gyrus at a coronal angle of ™14 degrees (AP ™2.0, ML ™1.86, DV ™1.24 mm). The cannula and the screws were then secured on the skull by acrylic materials. A headplate was fixed on the skull, and an opto-tetrode was implanted at the position targeting CA1 (AP ™2.0, ML ™1.5 mm) by gradually lowering the tetrode tip at the depth around 0.8 to 1.2 mm below the dura. The outer protective tube was put on the skull to isolate the opto-tetrode from acrylic materials^59^. A ground wire soldered on a skull screw was then connected to the female connector on the EIB.

### Seizure models and behavioral tests

KA-induced acute seizure model: KA (0.5 μg/μL, 0.6 μL) was unilaterally injected into the dentate gyrus (AP: ™2.0 mm, ML: ™1.25 mm, DV: ™1.6 mm) of mice using a syringe pump (KDS Legato 130, KD Scientific, USA) over a period of 10 min via the guide cannula (RWD Life Science, China). KA or saline was locally injected by inserting an injection cannula via the guide cannula pre-implanted with a 30-degree tilted angle. After recovery, seizure behavior of the mice was scored in accordance with the Racine scale at different scores: 1, sudden behaviour arrest; 2, facial jerking; 3, neck jerks; 4, clonic seizure or sitting; 5. tonic-clonic seizures (lying on belly); 6, tonic-clonic seizure (lying on side) or wild jumping; 7, tonic extension, possibly leading to respiratory arrest and death. Seizure scores 1–3 refer to focal seizures, and scores 4–5 refer to generalized seizures.

PTZ-induced seizure model: mice were intraperitoneally injected with PTZ (75 mg/kg, 54-95-5, TargetMol), and then evaluated for seizure severity and EEG monitoring in the next 90 min. In order to reduce the suffering or probability of death, all mice were euthanatized after the experiment. Seizure severity was scored according to the Racine scale.

Pilocarpine-induced seizure model: mice were intraperitoneally injected with lithium chloride (3 meq/kg, Sigma-Aldrich), and 20 hours later, mice were first administered methylatropine bromide (5 mg/kg, i.p.; Sigma-Aldrich) to suppress peripheral cholinergic activation by pilocarpine. Pilocarpine was administered 30 min later (350 mg/kg i.p.; 54-71-7, GlpBio), and mice were closely and continually monitored for behavioral indicators of seizures in the next 90 min. Seizure severity was scored in accordance with the Racine scale as mentioned above.

### Optogenetic stimulation and heating effect evaluation

Green laser light (561 nm wavelength) was delivered through a 200-mm diameter optical fiber connected to the LED laser (MSL-FN-561-S, Changchun New Industries Optoelectronics Tech., China) by a Master-8 commutator. The optical fiber was cut flat, and the laser power was adjusted to 3-20 mW. During behavioral test, optical fiber was secured to ensure no movement during the experiment. The stimulation parameters for intracranial illumination were: 10 ms light pulses at 10 Hz with either 10 s-on/10 s-off (short) or 2 min-on/1 min-off (long) cyclic light power around 6 mW (180 mW/mm^2^) measured at the outlet of the fiber tip; for transcranial optogenetics the light pulse width was increased from 10 ms to 50 ms, and the light power increase to around 15 mW.

In intracranial or transcranial optogenetic experiments, the BAT-12 Microprobe Thermometer (Physitemp Instruments, USA) was used to evaluate the heat effect induced by light stimulation. Briefly, mice were anesthetized and placed on a stereotaxic apparatus. After removing the fur of the head around the region of the interest, a burr hole was drilled on the exposed skull with a frame-mounted drill. The tip of the optical fiber (200 μm in diameter) was vertically placed either to the skull surface or at an angle to target light pulses to the brain region of interest. Thermocouple probes were inserted into the target brain region or placed at the brain surface for local temperature measurement.

### Tissue histology, staining, and imaging

#### Tissue sample preparation

Following injection with sodium pentobarbital (100 mg/kg; i.p.) to induce deep anesthesia, mice were transcardially perfused with ice-cold phosphate-buffered saline (PBS) followed by 4% paraformaldehyde (PFA) in PBS. Excised brains were then dissected out and post-fixed for 2-4 hours or overnight in 4% paraformaldehyde at 4°C on a shaker. Then, they were dehydrated in 30% sucrose solution and embedded in Tissue-Tek O.C.T. embedding compound (Sakura Finetek). Twenty-μm thickness brain slices attached on glass slides were prepared with a microtome (CM1950, Leica, Germany) and subsequently stored at - 80°C. Alternatively, 50-μm thickness brain sections were collected into 24-well plates and stored at ™80°C for free-floating immunostaining. For rat organotypic hippocampal slice cultures, sections were fixed in 4% PFA solution for 0.5 h and used directly for immunostaining.

Immunohistochemistry

In the immunostaining experiments, sections of mouse brains or rat hippocampal organotypic cultures were washed first in PBST solution (PBS buffer with 0.02% Triton X-100) for 30 min and blocked in the blocking solution (5% normal donkey serum in PBST solution) for 60 min at room temperature. They were incubated with primary antibodies overnight at 4 °C. Primary antibodies used in this study: rabbit anti-c-Fos (1:1000; 2250s, Cell signaling), guinea pig anti-RFP (1:500; GP-1080-50, Rockland), chicken anti-GFP (1:1000; ab13970, Abcam), rabbit anti-IBA-1 (1:500; ob-PRB029-01, Oasis biofarm), mouse anti-GFAP (1:500; MAB360, EMD Millipore), rabbit anti-DsRed (1:500; 632496, Clontech) and chicken anti-eGFP (to detect eYFP, 1:1000; A10262 Invitrogen) in the blocking solution. After washed three times in PBST solution for 10 min, sections were incubated with proper secondary antibodies: donkey anti-chicken IgY/mouse IgG Alexa Fluor 488, donkey anti-guinea pig/rabbit IgG Cy3 (1:500; Jackson ImmunoResearch), goat anti-rabbit IgG Alexa Fluor 647 (1:1000; A27040, Invitrogen) or goat anti-chicken IgY Alexa Fluor 488 (1:1000; A11039, Invitrogen) spiked with DAPI (ab104139, Abcam) for 2 h at room temperature. Then sections were washed three times in PBST solution for 10 min and mounted with the Fluoroshield mounting medium (F6182, Sigma-Aldrich). Images were acquired with a confocal microscope (AiryScan 900, Zeiss, Germany).

#### *In situ* hybridization

*In situ* hybridization experiments were performed following the method previously described^60^. Fluorescein- or DIG-labeled mCaMKIIα probes used for hybridization on 50 μm free-floating cryosections. Hybridization was performed overnight at 65°C. Sections were washed at 60°C twice in 2x SSC (saline sodium citrate) solution with 50% formamide and 0.1% N-lauroylsarcosine, treated with 20 μg/ml RNase A for 15 min at 37°C, then washed twice in 2x SSC solution with 0.1% N-lauroylsarcosine at 37°C for 20 min and in 0.2x SSC solution with 0.1% N-lauroylsarcosine twice at 37°C for 20 min. Sections were blocked in MABT solution containing 10% goat serum and 1% blocking reagent (11096176001, Roche) for 1 h at room temperature. Then they were incubated with anti-Fluorescein-POD (11426346910, Roche) or anti-DIG-POD antibody (11207733910, Roche) for 1 h at room temperature. After a washing step, signal amplification TSA staining was performed using Fluorescein or Cy3 TSA Fluorescence System Kit (K1050- 100-300, APExBIO). Images were acquired using a fluorescence microscope (Axio Imager.M2, Zeiss, Germany) or a confocal microscope (LSM 710; Zeiss, Germany) and quantified using Image J software (v1.8.0).

### Statistics and reproducibility

Data analyses were performed using GraphPad Prism (Version 8.4.3 or 9.0.0, San Diego, CA) and MATLAB (Version 2018b). Data are presented as mean ± standard error of the mean (SEM) or median and interquartile range as indicated. Statistical details for specific experiments - including exact number of repeats (n), p values, and statistical tests are specified in figure legends. Where representative images are shown, each experiment was repeated at least three times independently with similar results. Data were analyzed by one-way or two-way ANOVA followed by *post hoc* Dunnett’s, Tukey’s or Sidak’s test for multiple comparisons. In cases where normality/equal variance failed, a non-parametric Friedmann test was used instead of ANOVA. When comparing two conditions an unpaired Mann-Whitney test (data non-normal), unpaired two-tailed t-test (data normal, equal variance), or paired t-test for statistical significance when appropriate. For Kaplan-Meier plots showing survival rate, a log-rank (Mantel-Cox) test was used. Significance levels are denoted as \**P* < 0.05, \*\**P* < 0.01, \*\*\**P* < 0.001, \*\*\*\**P* < 0.0001; ns, not significant. The details of statistical tests are described in the figure legends.

## Acknowledgments

We thank the Animal Facility for animal maintenance and Core Research Facility for imaging support at the Southern University of Science and Technology. We appreciate Prof. Duanqing Pei and Dr. Mingfeng Zhang sharing the structure of KCR1. This study was supported by grants from the Shenzhen Innovation Committee of Science and Technology-Shenzhen Key Laboratory of Gene Transcription and Systems Biology (ZDSYS20200811144002008 to K.S.) and Shenzhen-Hong Kong Institute of Brain Science-Shenzhen Fundamental Research Institutions (2023SHIBS0002 to K.S. and S.T.H.). STH is also supported by the Guangdong Innovation Platform of Translational Research for Cerebrovascular Diseases and SUSTech-UQ Joint Centre for Neuroscience and Neural Engineering (CNNE).

## Author contributions

K.S. and X.D. conceived the project, designed the experiments and wrote the manuscript. X.D. and Y.W. performed the seizure induction, fiber photometry recording, and behavior tests *in vivo* as well as the immunohistology experiments. S.G. G.N. and C.Z. engineered the KCR1 variants including iKCR and hsKCR and characterized them in oocytes. C.G., S.O. and O.M.C. compared the electrophysiological properties of KCR1, iKCR and hsKCR on organotypical brain slices. S.H., J.J. and X.L. performed the electrophysiological recordings on acute brain slices. X.C., C.W. and Z.X. performed the LFP recordings. Z.L. and Y.L. helped with *in situ* hybridization. All the authors read and revised the manuscript.Competing interests K.S., X.D. C.Z. and Y.W. have filed a patent application related to this work. The remaining authors declare no competing interests.

## Notes

### Competing Interest Statement

The authors have declared no competing interest.

