## supplementary materials for "Suppression of epileptic seizures by transcranial activation of K^+^-selective channelrhodopsin"

### Extended Data

Extended Data Figure 1

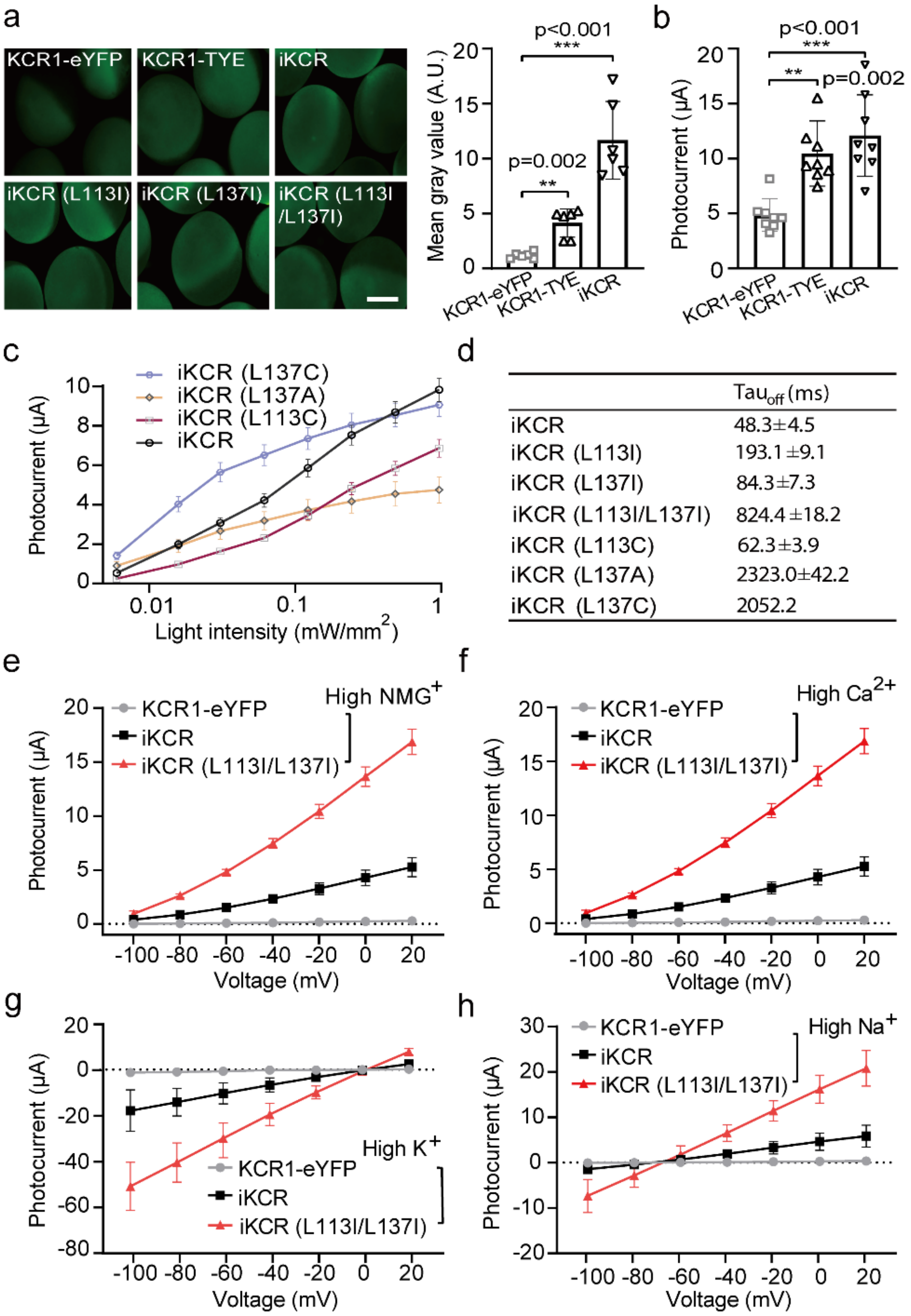


**Extended Data Fig 1. Expression and ion selectivity of KCR1 variants in *Xenopus* oocytes. a,** Expression of KCR1 mutants in *Xenopus* Oocytes. Scale bar = 0.5 mm. Expression levels were quantified according to the grey value of the cell images. **b,** Photocurrent amplitudes of KCR1 and optimized KCR1 variants measured by two-electrode voltage clamp at a holding potential of -40 mV in response to 1 mW/mm$^2$, 530 nm, 1 s light pulses. The bath solution was ORi with 2 mM $BaCl_2$, n=6 oocytes. **c,** Light sensitivity of iKCR and its point mutants, n=7-9 oocytes; light stimulation condition 530 nm, 1 s. **d,** Closing kinetics using the same stimulation parameters as in **(b),** n=6 oocytes. **e-h,** I-V relationship of KCR1-eYFP, iKCR and iKCR (L113I/L137I) in the indicated solutions, n=6 oocytes. The contents of the solutions are shown in **Extended Data Table 1**. Statistical analysis was performed by unpaired Mann-Whitney test. Error bars are standard errors of the mean (SEM).

Extended Data Figure 2

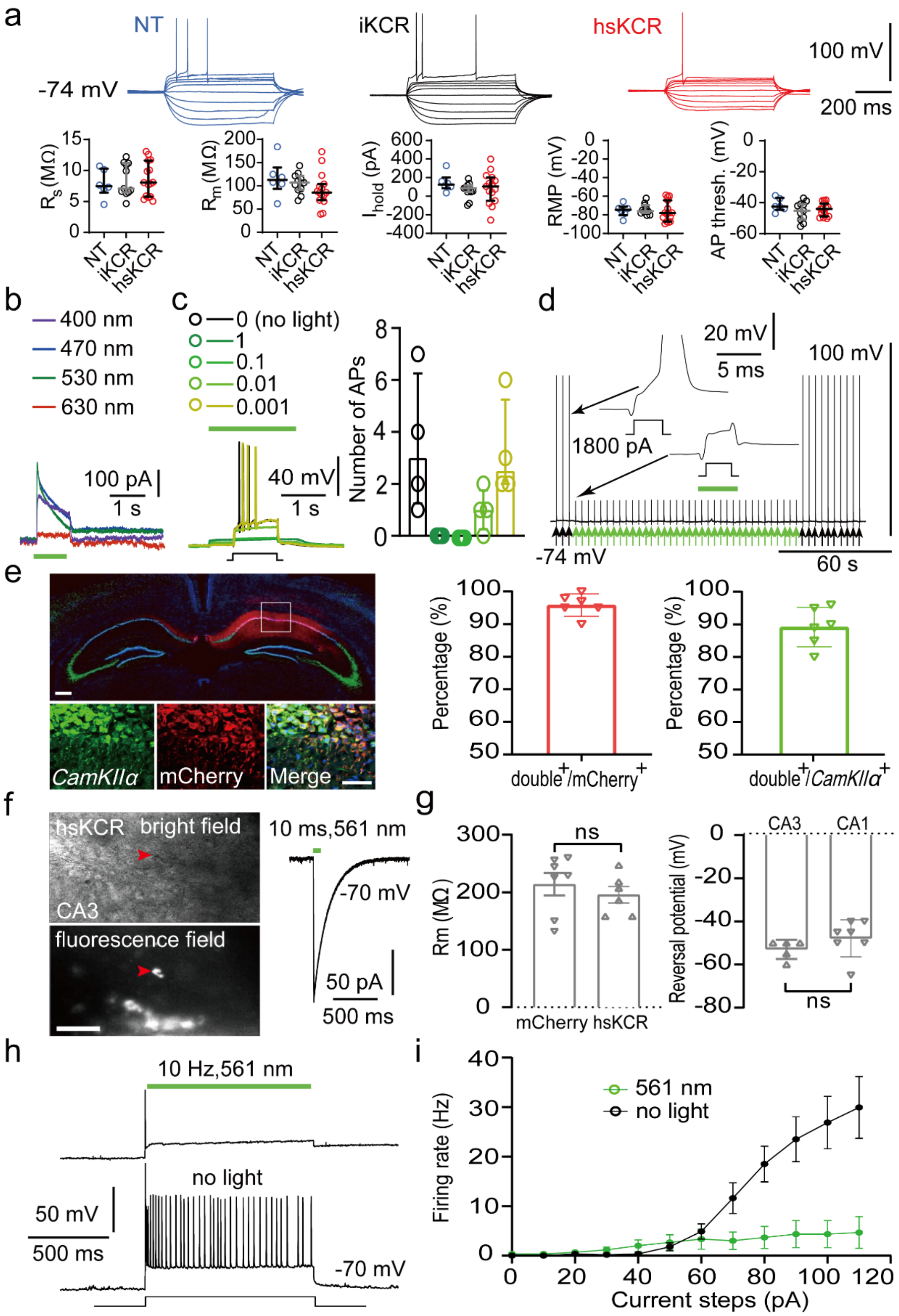


**Extended Data Fig 2: Functional characterization of iKCR and hsKCR in rodent hippocampal neurons. a,** Membrane potential responses to current injection steps from -400 pA to 400 pA of exemplary non-transduced (NT), iKCR-expressing and hsKCR-expressing rat CA3 neurons in organotypic slice cultures. Series resistance ($R_s$), membrane resistance ($R_m$), holding current ($I_{hold}$) when voltage clamped at -74 mV, resting membrane potential (RMP) and the threshold for action potential firing (AP thresh.) were not different among NT, iKCR (DNA electroporated at 10 ng/µl) and hsKCR (DNA electroporated at 1 ng/µl) expressing CA3 neurons (median and interquartile range). **b,** Representative photocurrents recorded at -60 mV in an iKCR-expressing CA3 pyramidal neuron activated by the light with different wavelengths. The green bar indicates 1 s, 1 mW/mm$^2$ light illumination. **c,** Exemplary current-clamp recordings of action potentials from an iKCR-expressing CA3 neuron during 1 s current injection (black step, left panel). The green bar indicates 1 s application of 530 nm light of various intensities. Statistical analysis of the relationship between the number of action potentials and light intensities (median and interquartile range, p=0.0008, Friedmann test, n=4 cells). **d,** Action potentials of an iKCR-expressing neuron in response to 3 ms 1800 pA current pulses every 5 s (arrows below traces, black steps in inserts). Starting from the $4^{th}$ pulse, green light pulses were triggered 1 ms before the current steps (green arrows/green bar, 5 ms, 530 nm, 10 $mW/mm^2$). **e,** Representative images of neurons expressing mCherry$^+$ (immunostaining) or CaMKIIα$^+$ (*in situ* hybridization) in hippocampal CA1 region with a unilateral injection of AAV2/9-mCaMKIIα-hsKCR-mCherry (left panel, scale bars, 100 µm). Quantification of the coexpression ratios between mCherry and CaMKIIα (right panel, n=6 mice). **f,** Bright-field and fluorescence images of an hsKCR-expressing neuron (red arrow) in an acute mouse hippocampus CA3 slice and its exemplary photocurrent. Scale bar: 100 µM. **g,** Quantification of the hsKCR reversal potential and membrane resistance. Membrane potentials were not corrected for LJP (liquid junction potentials), which was measured experimentally around - 13 mV. Unpaired *t*-test (n=5-7 cells from 3 mice). **h** and **i**, Activation of hsKCR by 10 Hz green light significantly suppressed the firing of hsKCR-expressing CA3 pyramidal neurons. Representative traces (**h**) and statistical analysis (**i**, n=6-8 cells from 3 mice).

Extended Data Figure 3

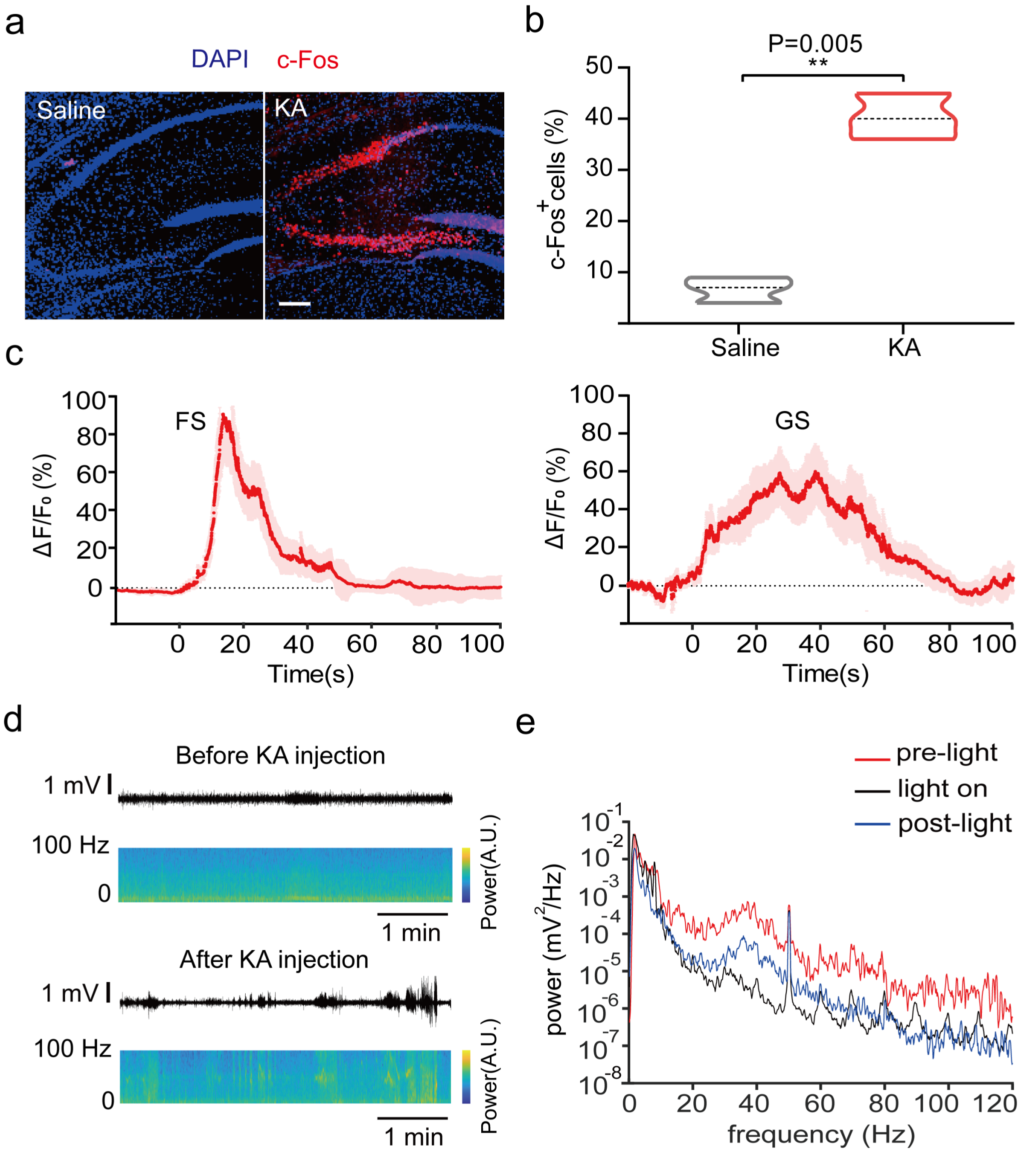


**Extended Data Fig 3. Activation of hsKCR significantly inhibited epileptic seizure. a**, c-Fos immunostaining images of mouse hippocampi treated with saline or KA (0.5 μg/μl, 600 nl applied via a pre-implanted cannula). **b,** Quantification of c-Fos-expressing cells in the hippocampi of saline or KA treated mice, unpaired *t*-test, n=3 mice, scale bar: 100 μm. **c,** Transient $Ca^{2+}$ signals of CaMKIIα$^+$ hippocampal CA3 neurons during FS and GS induced by KA injection (n=14 trails from 7 mice). **d,** Typical LFP recording traces from hsKCR-expressing mice before and after KA injection. **e,** Analysis of power spectral density during light (light gray), before light (red), and after light (blue) illumination.

Extended Data Figure 4

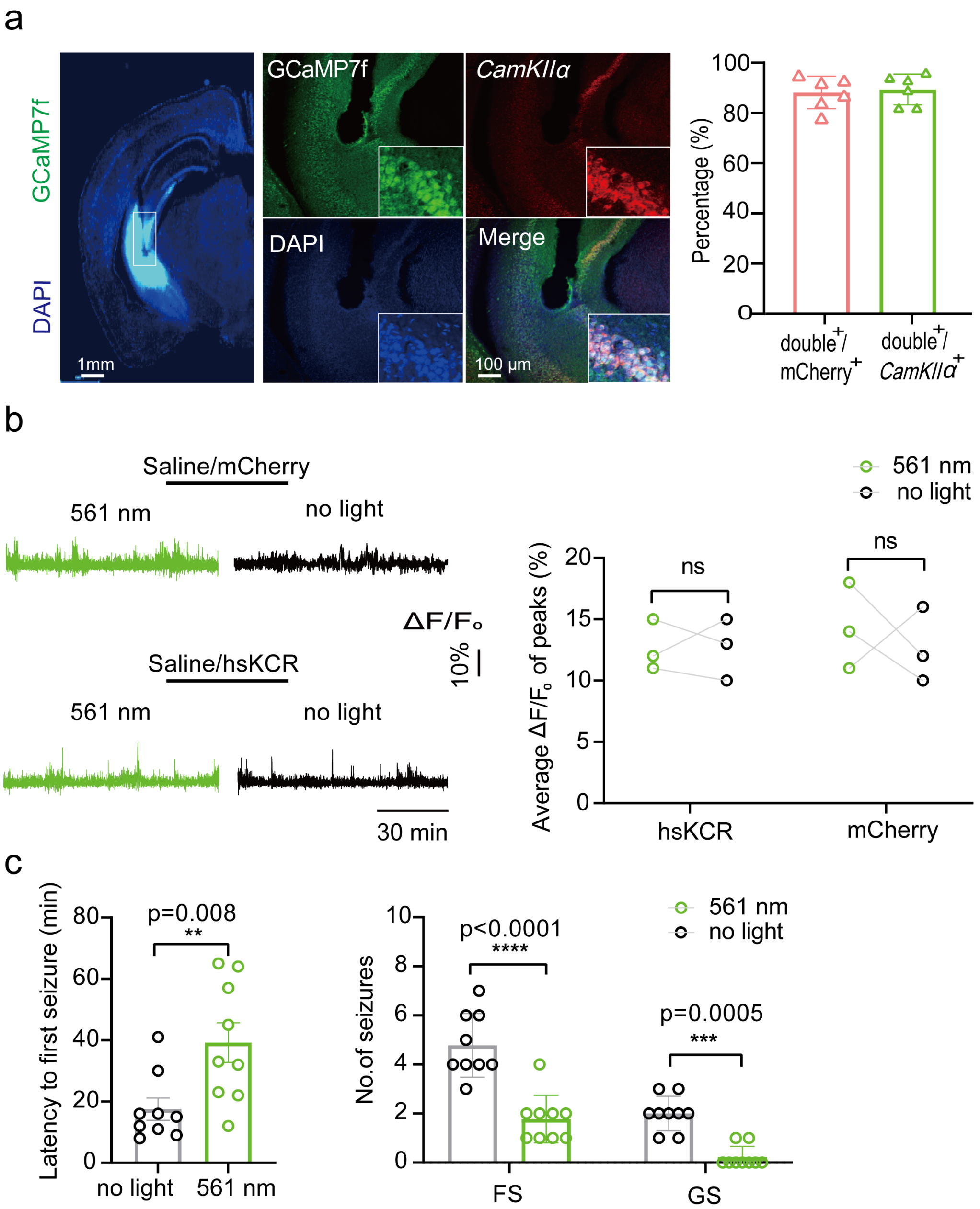


**Extended Data Fig 4. Specific expression patterns of GCaMP7f in CaMKIIα⁺ pyramidal neuron. a,** Fluorescent hippocampal images of GCaMP7f (detected by a GFP antibody) and CaMKIIα mRNA (detected by *in situ* hybridization) staining in hippocampal CA3 area of a mouse unilaterally injected with AAV2/9-mCaMKIIα-GCaMP7f and the quantification of the co-expression ratios of GCaMP7f and *CaMKIIα* in pyramidal neurons. n=6 mice; scale bars, 1 mm (left) or 100 μm (right). **b,** Representative traces of $Ca^{2+}$ recording in mCherry- or hsKCR-expressing mice with or without 561 nm light illumination after saline injection. $Ca^{2+}$ change was indicated as $\Delta F/F_0$ (%). The statistical analysis of $Ca^{2+}$ changes was presented in the right panel. n=3 mice. **c,** Statistical analysis of behavioral seizures indicated as latency to first seizure (left, Mann-Whitney test) and the numbers of FS or GS (right, two-way ANOVA with Šídák's multiple comparison test) in the hsKCR-expressing mice with or without 561 nm light illumination after KA treatment. n=9 mice. Irradiation condition: cyclic 561 nm laser light (10 ms light pulse at 10 Hz for 2 min, and then 1 min no light) with light power around 6 mW (180 mW/mm$^2$) measured at the fiber tip.

Extended Data Figure 5

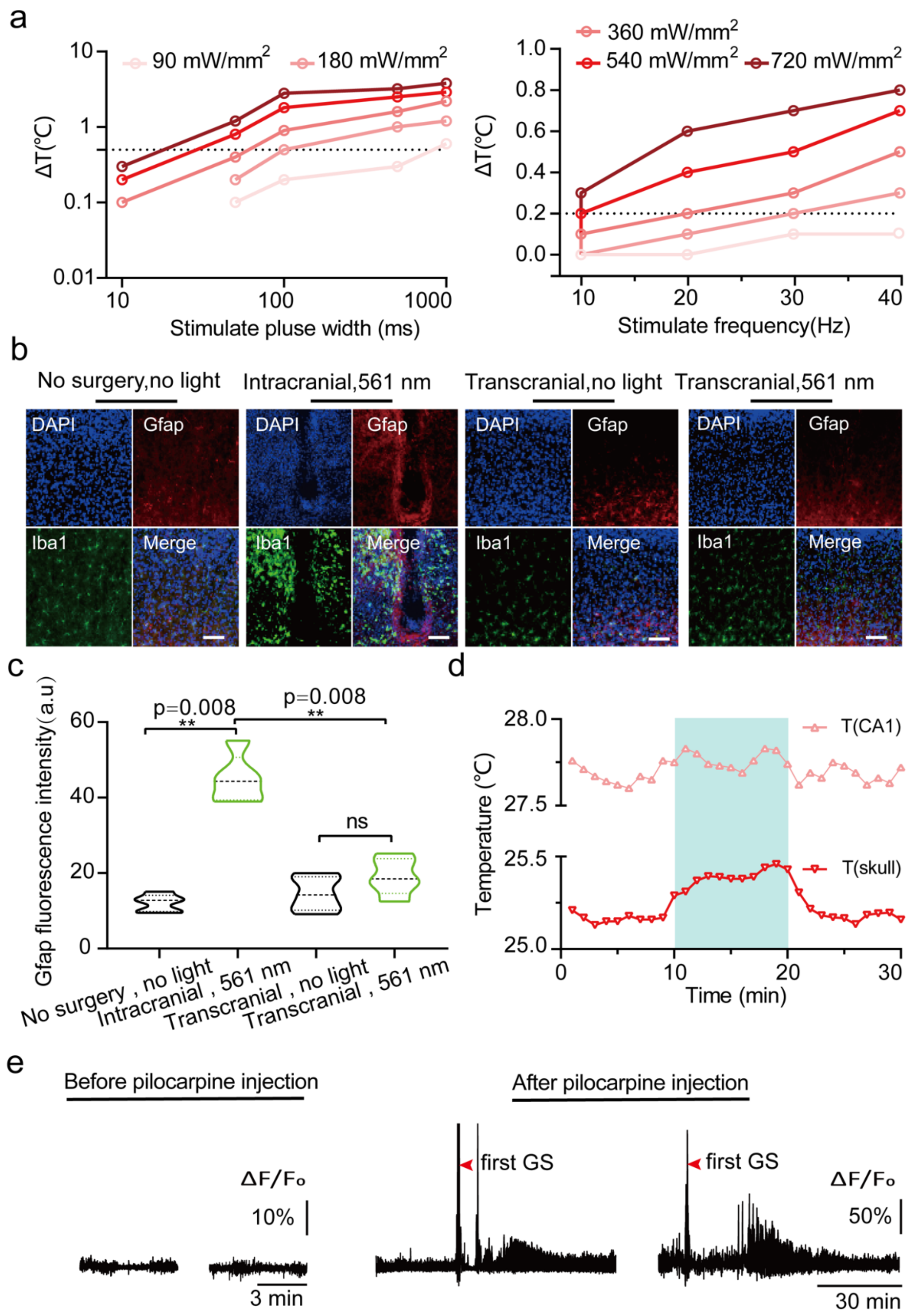


**Extended Data Fig 5. Optimization of light irradiating conditions for transcranial optogenetic application. a,** Temperature changes associated with a 561 nm laser illumination at different pulse widths, frequencies and powers of the green light, n=3 mice. **b,** Comparison of Iba-1 and Gfap expression levels from different mice without surgery under no light condition, intracranial optogenetic stimulation and transcranial optogenetics with or without light. Scale bars, 100 μm. **c,** Quantification of Gfap fluorescence intensity in the regions indicated by the black boxes shown in **Fig 5a**. One-way ANOVA with Tukey's multiple comparisons test, n=5 mice. **d,** Measurement of temperature change during transcranial optogenetic stimulation at the CA1 region and skull surface**. e,** Representative $Ca^{2+}$ activity before and after pilocarpine injection in hsKCR-expressing mice.

Extended Data Figure 6

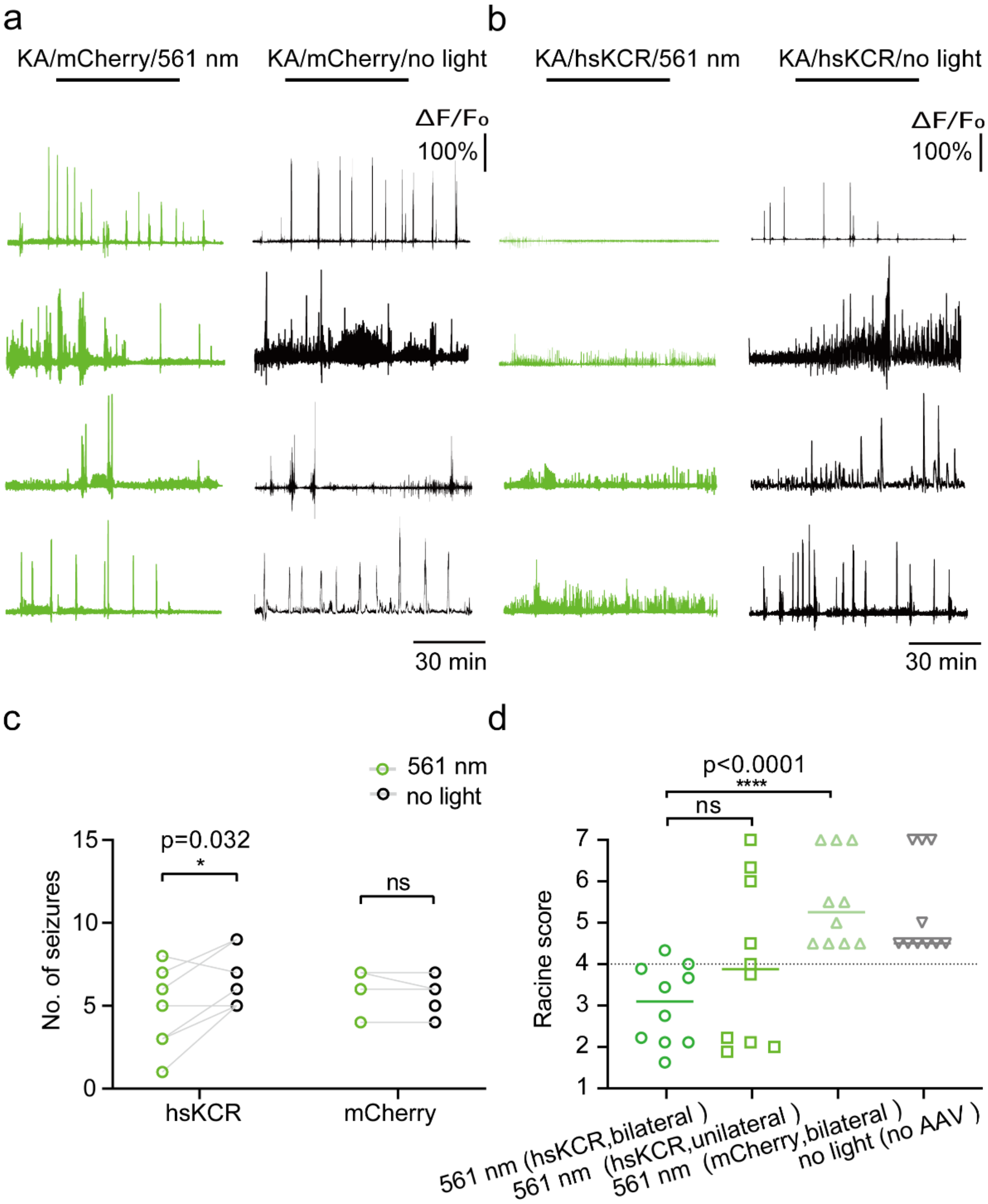


**Extended Data Fig 6. Bilateral transcranial optogenetics significant inhibited behavioral seizure in epileptic mice.** **a** and **b,** Representative $Ca^{2+}$ recording traces from mCherry (**a**) or hsKCR-mCherry (**b**) expressing mice with 561 nm green light (left) or without light (right) illumination during 1.5 h KA kindling period; light stimulation condition: 561 nm laser, 50 ms light pulse at 10 Hz with light power around 15 mW (450 mW/mm$^2$) measured at the optical outlet. $Ca^{2+}$ change was indicated as $\Delta F/F_0$. **c,** Quantification of behavioral seizure indicated by number of seizures in mCherry or hsKCR expressing mice with or without transcranial light delivery 1.5 h after KA treatment, mCherry mice n=4, hsKCR mice n=7. Multiple paired *t*-test, ns, no significant, *P < 0.05. **d,** Racine Score of PTZ-treated mice under different optogenetic conditions. One-way ANOVA with *post hoc* Dunnett's test. n=10.

**Extended Data Table 1. Solutions for the two-electrode voltage-clamp measurements**

Solution contents (in mM)

| | NaCl | KCl | NMG | $MgCl_2$ | $CaCl_2$ | pH |
|---|---|---|---|---|---|---|
| Ringer solution | 110 | 5 | | 1 | | 7.6 |
| High Na | 115 | | | 1 | 2 | 7.5 |
| High K | | 115 | | 1 | 2 | 7.5 |
| NMG-Cl | | | 115 | 1 | 2 | 7.5 |
| 20 mM $CaCl_2$ | | | 86 | 1 | 20 | 7.5 |
